# Deciphering Peptide-Protein Interactions via Composition-Based Prediction: A Case Study with Survivin/BIRC5

**DOI:** 10.1101/2024.02.12.579763

**Authors:** Atsarina Larasati Anindya, Torbjörn Nur Olsson, Maja Jensen, Maria-Jose Garcia-Bonete, Sally P. Wheatley, Maria I. Bokarewa, Stefano A. Mezzasalma, Gergely Katona

## Abstract

In the realm of atomic physics and chemistry, composition emerges as the most powerful means of describing matter. Mendeleev’s periodic table and chemical formulas, while not entirely free from ambiguities, provide robust approximations for comprehending the properties of atoms, chemicals, and their collective behaviours, which stem from the dynamic interplay of their constituents.

Our study illustrates that protein-protein interactions follow a similar paradigm, wherein the composition of peptides plays a pivotal role in predicting their interactions with the protein survivin, using an elegantly simple model. An analysis of these predictions within the context of the human proteome not only illuminates the known cellular locations of survivin and its interaction partners, but also introduces novel insights into biological functionality. It becomes evident that an electrostatic- and primary structure-based descriptions fall short in predictive power, leading us to speculate that protein interactions are orchestrated by the collective dynamics of functional groups.

## INTRODUCTION

Most biological processes are directed by protein-protein interactions (PPI). These interactions are associated with a network of non-covalent bonds in a way that are not completely understood, and their deficiencies are often related with disease progression, i.e. aggregation and amyloid formation.

Early studies readily account interactions between proteins with opposite charges, and that increasing the solution ionic strength reverses the interaction^1,2^. Short-range forces such as hydrogen bonding and van der Waals’ were also acknowledged to contribute^3^. After insulin was successfully sequenced in 1951^4^, the growing number of sequenced proteins brought forth a paradigm to treat the numerous possible combinations of protein primary amino acid sequence as the sole key to determine protein structure, function, and, by extension, how it recognize interaction partners, often oversimplifying the physicochemical properties of the amino acids embodying the biological message.

Each amino acid indeed has unique properties that ultimately contribute to the overall electrostatic and polar/apolar nature of the protein (charge density and distribution, net charge, and hydrophobicity). However, the physical description based on complementary charge of two partners and similarity in hydrophobicity are not regarded to be sufficiently detailed to direct specific interactions.

At the same time, research on identifying protein binding partners and determining binding affinity keeps rising with the help of high-throughput techniques like two hybrid yeast^5,6^, phage display^7^, bimolecular fluorescence complementation^8^, and peptide microarrays^9,10^. Our understanding of these interactions is not happening with the same pace and generally explained by detailed short-range physical interactions. Black box machine learning of cognate partners runs the risk of merely transforming existing data into yet another (overfit) model representation.

Peptide microarrays, which has its roots in antigen-antibody reaction determination method^11^, is an effective tool in studying PPI due to its high-throughput nature. Simultaneous determination of PPI is performed by exposing multiple antibody spots, arranged on a solid surface, to antigens. In peptide microarrays, the target protein is often divided into short peptide segments and orderly arranged on a solid surface, before being exposed fluorescence-labelled ligand proteins or antibody in solution. The detected fluorescence intensity at each peptide spot correlates to the binding strength of that definite peptide to the ligand protein^12^, making mapping binding and non-binding regions in a protein possible, as we have previously shown with the mapping of Polycomb Repressor Complex 2 (PRC2) binding regions to survivin^13^.

Rapid accumulation of PPI experimental data from peptide microarray becomes a valuable factor to formulate patterns to identify and understand the driving forces behind. A major challenge to derive prediction tools comes from inconsistencies on defining the cut-off distance of non-covalent interactions^14^, which are modelled based on amino acid physicochemical and biochemical properties; AAindex is a database example containing various indices representing these properties^15^.

Protein association is thought to start with encounter complex formation mediated by long-range electrostatic forces resulting in a favourable (exothermic) enthalpy change^16^, which can likely be predicted by statistical coupling analysis^17^. Encounter complex formation implies an entropic loss from reduced conformational, rotational, and translational degrees of freedom before definite hydrogen and ionic bonds are formed, leading to structural rearrangements and eventual desolvation of the interacting surfaces^18–20^.

Certain characteristics are connected to protein interfaces: stable interactions have larger buried surface areas and hydrophobic surfaces than transient ones^21^. Stable interactions also have more conserved and tryptophane, tyrosine, and arginine residues, where most of the binding energy is concentrated^22,23^. The interface centre of stable interactions is usually composed of hydrophobic and aromatic residues, while peripheral regions are often crowded with polar and charged residues. Interfaces with stronger affinities tend to have high modularity involving large, complex interfacial sub-regions, suggesting that individual pairwise interacting residues have additive effects^24,25^. Side-chain interactions are also prominent in PPIs, whereas they only contribute to 36% of the internal bonds stabilizing a protein^26^. There is also a significant compositional difference between transient and permanent interactions (i.e. in quaternary structures or complexes)^27^.

Despite knowing conserved residues forming PPIs, binding affinity prediction based solely on amino acid sequence have not been unambiguous, as diverse sequences can have similar binding affinities^28^. However, amino acid sequence data remains the most common feature for inferring structure, and by extension, protein activity. Multiple Sequence Alignment combined with structural template-trained model is the current state-of-the-art strategy employed by well-known prediction tools, such as AlphaFold Multimer^29^, AlphaPullDown^30^, and RoseTTAFold^31^. These tools have limitations on proteins with no solved homologous structure, a common problem with inherently disordered proteins (IDPs) and transient/weak binding contact interfaces.

Methods using feature extraction from sequence alignment is often inspired from natural language processing methods, such as masked language^32^, attention network, and long short-term memory (LSTM) models^31,33–35^. The last two are currently developed to counter the problem of long-range communications between residues, which is found in classical recurrent neural network. Cadet et al recently developed a model relying on the Fourier transform of numerical-encoded sequences, creating a unique pattern from residue frequency for each protein ^36^. This study used a training dataset of single point mutations of epoxide hydrolase from *Aspergillus niger* and achieved 81% accuracy on validation dataset to predict epistatic interactions^36^. On the other hand, binding site predictions from the joint analysis of solvation potentials, amino acid composition, conservation, electrostatics, and hydrophobicity seems to yield more promising results^37–42^.

Chemical reactions in dynamic equilibrium, including PPIs, are assumed to depend on the law of mass action, where the product to reactant ratio is constant for given temperature. The amount of protein complex depends on interacting partner concentrations and their binding affinity. The scarcity of explicitly repulsive PPI observations points to a prevalence of attractive interactions among proteins or to an observational bias focusing on attractive interactions.

Considering the large size and complexity of individual proteins, and the diversity among proteins, it is of practical interest to shift the focus from the concentration of polypeptide chains and protein complexes to whether the concentration of simpler entities composing proteins plays a pivotal role in guiding PPIs. Here, we use survivin, a small protein (16.5 kDa) involved in apoptosis, cell division, and epigenetic regulation for transcription^13^, as the target protein model and its binding partners to illustrate how protein composition is a predictive factor for biological interactions. We validate our assertion by referring to extensive experimental data available in the literature about survivin interaction partners and the role of survivin in various biological functions. Finally, we discuss the effectiveness of composition in determining protein function versus the collective dynamics of proteins.

## RESULTS AND DISCUSSION

### PPI classification based on atom type composition and sequence

This study used a peptide microarray dataset containing fluorescence intensity from survivin interactions to 36 known and potential binding partners^13^, such as Polycomb Repressive Complex 2 (PRC2) subunits^43^ and Chromosomal Passenger Complex (CPC). The peptides were then represented by counting various chemical groups they contain as features, encompassing the standard main chain atoms (MC), methylene group from glycine Cα atom (CA-Gly), side chain atoms (carboxyl, amide, phenolic hydroxyl, sulfhydryl groups, imidazole and indole rings) among others. A comprehensive list and explanation on feature development can be found in the Methods section, with cues to appropriate Supplementary Figures and Tables. Briefly, C-Coord1 model as a starting step contains 17 features (Table S1), C-Coord2 with 15 features (Table S2), and C-Coord3 with 13 features (Table S3). C-Pos model characterizes the distance of side chain carbon atoms from the main chain (Table S4), AA-Letter model (Table S5) simply substitutes amino acids with equal weight values, and the chemical formula model counts hydrogen, carbon, nitrogen, oxygen, and sulfur atoms (Table S6). Electrostatic and hydrophobic properties are represented in Pos/Neg/HyPho model as binary properties of individual amino acids (Table S7) and NetChrg/SumHyPho models representing the properties of the entire peptide (Table S8)^44^. We also compared our approach with Fourier Transform patterns from AAindex numerical value representation of amino acids^36^.(Seq, Table S9) The real part of the complex term yields a peptide spectrum (FT-Seq). We also used equal weight value representation of amino acids in alphabetical order (from A=10 to W=200) (UniSeq).

Figure 1 presents the performance indicators of various approaches, using a very simple multilayer perceptron with two intermediate layer perceptrons. As previously reported^13^, non-binders are slightly more common compared to binders. Models with more limited features exhibit suboptimal performance, with accuracy and F1 scores for binders falling within the range of 65—74 % and 0—62%, respectively. The FT-Seq, FT-UniSeq feature sets, and the simplest physicochemical descriptor (NetChrg/SumHyPho) similarly exhibit underperformance. It is intriguing to observe that the elemental composition provides a more effective description compared to features based on electrostatic and hydrophobicity characteristics of the peptides. Conversely, symbolic AA-Letter description exhibits lower performance when employed in a simple multilayer perceptron model. We speculate that the AA-Letter feature set lacks a direct alignment with fundamental chemical principles, i.e. it fails to consider the varying numbers of atoms among different amino acids and does not reflect any similarities between them. More sophisticated models may learn these aspects with sufficient data (as demonstrated in the embedding example later), yet the C-Pos, C-Coord1, and C-Coord2 notably align better with intuitive understanding of amino acid features with limited data.

**Figure 1.**
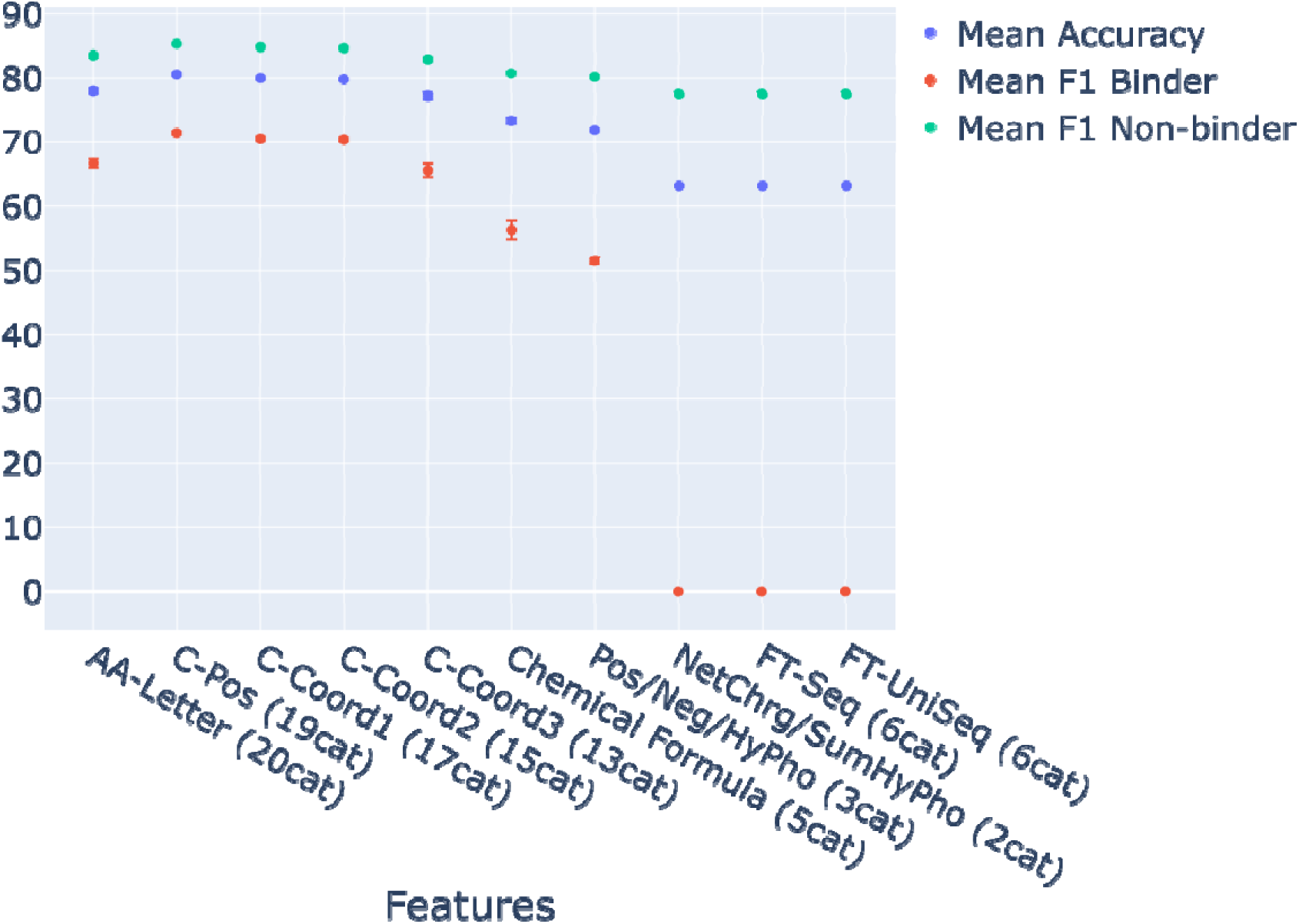
Performance evaluation of a multilayer perceptron classification with a single intermediate layer consisting of two perceptrons. Different sets of features describing the peptides on the peptide microarray were compared. Training was carried out using 50% of the data, repeated 100 times with varying randomly selected subsets, and testing was performed on the remaining half of the dataset.

Simplification resulted in similar performances when the 15-feature model (C-Coord2) is compared with a 17-feature model (C-Coord1). However, overall performance declined with reduction to 13 (C-Coord3) features, suggesting the crucial role of differentiating oxygen atoms in carboxyl and carbonyl groups and nitrogen atoms in amino and amide groups in prediction. As C-Pos and C-Coord1 only differ in the labeling of aliphatic carbon atoms, the slightly better performance of the C-Pos model implies the importance of aliphatic carbon atoms in biomolecular recognition, an area where current awareness is limited.

The C-Coord2 model demonstrated an average accuracy of 79.8% (Table 1) compared to the FT-Seq with 63.2% accuracy by only predicting non-interacting peptides. Specifically, the F1 scores for detecting non-binders and binders were 85% and 71% for the C-Coord2 model, whereas the FT-Seq approach failed completely to detect binders. The performance indicators show that the model is biased towards non-binding peptides, which might be explained by the slightly imbalanced dataset, comprised of 60% non-binders.

**Table 1.** Performance indicators of the survivin binding peptide predictions. Uncertainty is represented by the standard error of the mean.

|  | Precision (%) |  | Recall (%) |  | F1-score (%) |  | Accuracy (%) |
| --- | --- | --- | --- | --- | --- | --- | --- |
|  | NB | B | NB | B | NB | B |  |
| C-Coord2 | 81.6 ± 0.3 | 76.0 ± 0.4 | 87.9 ± 0.3 | 65.8 ± 0.5 | 84.6 ± 0.2 | 70.5 ± 0.3 | 79.8 ± 0.2 |
| C-Coord2<br>every 3<sup>rd</sup><br>peptide | **80.6 ± 0.6** | 69.9 ± 1.4 | 83.4 ± 1.2 | **65.5 ± 1.2** | 81.9 ± 0.6 | 67.4 ± 0.8 | 76.8 ± 0.7 |
| C-Pos | 81.9 ± 0.3 | 77.7 ± 0.5 | 89.0 ± 0.3 | 66.1 ± 0.6 | 85.3 ± 0.1 | 71.4 ± 0.3 | 80.6 ± 0.2 |
| C-Pos<br>every 3<sup>rd</sup><br>peptide | 80.4 ± 0.6 | **75.3 ± 0.9** | **87.6 ± 0.7** | 63.6 ± 1.3 | **83.8 ± 0.2** | **68.8 ± 0.5** | **78.7 ± 0.3** |
| Seq | 64.3 ± 0.3 | 60.2 ± 5.0 | 98.1 ± 0.4 | 5.6 ± 1.0 | 77.7 ± 0.2 | 10.0 ± 1.8 | 64.2 ± 0.3 |
| UniSeq | 63.7 ± 0.2 | 47.8 ± 1.5 | 96.6 ± 0.4 | 5.3 ± 0.5 | 76.8 ± 0.2 | 9.4 ± 0.9 | 63.1 ± 0.2 |
| FT-Seq | 63.2 ± 0.2 | 0.0 ± 0.0 | 100.0 ± 0.0 | 0.0 ± 0.0 | 77.4 ± 0.2 | 0.0 ± 0.0 | 63.2 ± 0.2 |

**Table 2.** Confusion matrix for the prediction of survivin-microarray peptide interactions using the LSTM model using the cross-validation protocol. Uncertainty is represented by the standard error of the mean.

|  |  | Predicted label | Every peptide |  | Predicted label | Every Third Peptide |
| --- | --- | --- | --- | --- | --- | --- |
|  |  | Non-interacting | Interacting |  | Non-interacting | Interacting |
| Correct label | Non-interacting | 1218±4 | 159±3 | Non-interacting | 379±2 | 77±2 |
|  | Interacting | 228±4 | 1093±4 | Interacting | 110±3 | 333±3 |
| Precision (%) |  | 84.3±0.2 | 87.4±0.2 |  | 77.8±0.4 | 81.6±0.4 |
| Recall (%) |  | 88.5±0.2 | 82.7±0.3 |  | 83.2±0.5 | 75.2±0.6 |
| F1-score (%) |  | 86.3±0.1 | 85.0±0.1 |  | 80.2±0.2 | 78.0±0.3 |
| Accuracy (%) |  | 85.7±0.1 |  |  | 79.2±0.2 |  |

### The impact of network complexity on the prediction accuracy and applicability

Next, we investigated how the complexity of the multilayer perceptron impacts prediction performance with different feature sets. To maintain simplicity, we employed a single intermediate layer of perceptrons and explored changing the number of hidden layer perceptrons, ranging from the default 100 down to as few as 1. We also evaluated the performance without relying on a distinct test set to assess the model susceptibility in making predictions based on insignificant details and noise within the training set (overfitting).

By decreasing model complexity, composition-based models not only increased overall accuracy but also significantly enhanced the more challenging detection of binders, as demonstrated by the F1 Positive score (Figures 2A and 2B). The optimal configuration, with peak accuracy and F1 scores, emerges with a two-perceptron intermediate layer. Remarkably, there is minimal disparity between the performance of the model with and without a test set, indicating a well-balanced accuracy and generalization. Notably, there exists a substantial non-monotonic variation in F1 performance, even with minor alterations in the number of perceptrons, suggesting that specific logical configurations are more advantageous than others when evaluating the connectivity between perceptrons. The subpar performance of the single perceptron model refutes the argument to dismiss amino acid and atom type diversity for precise binder detection.

**Figure 2.**
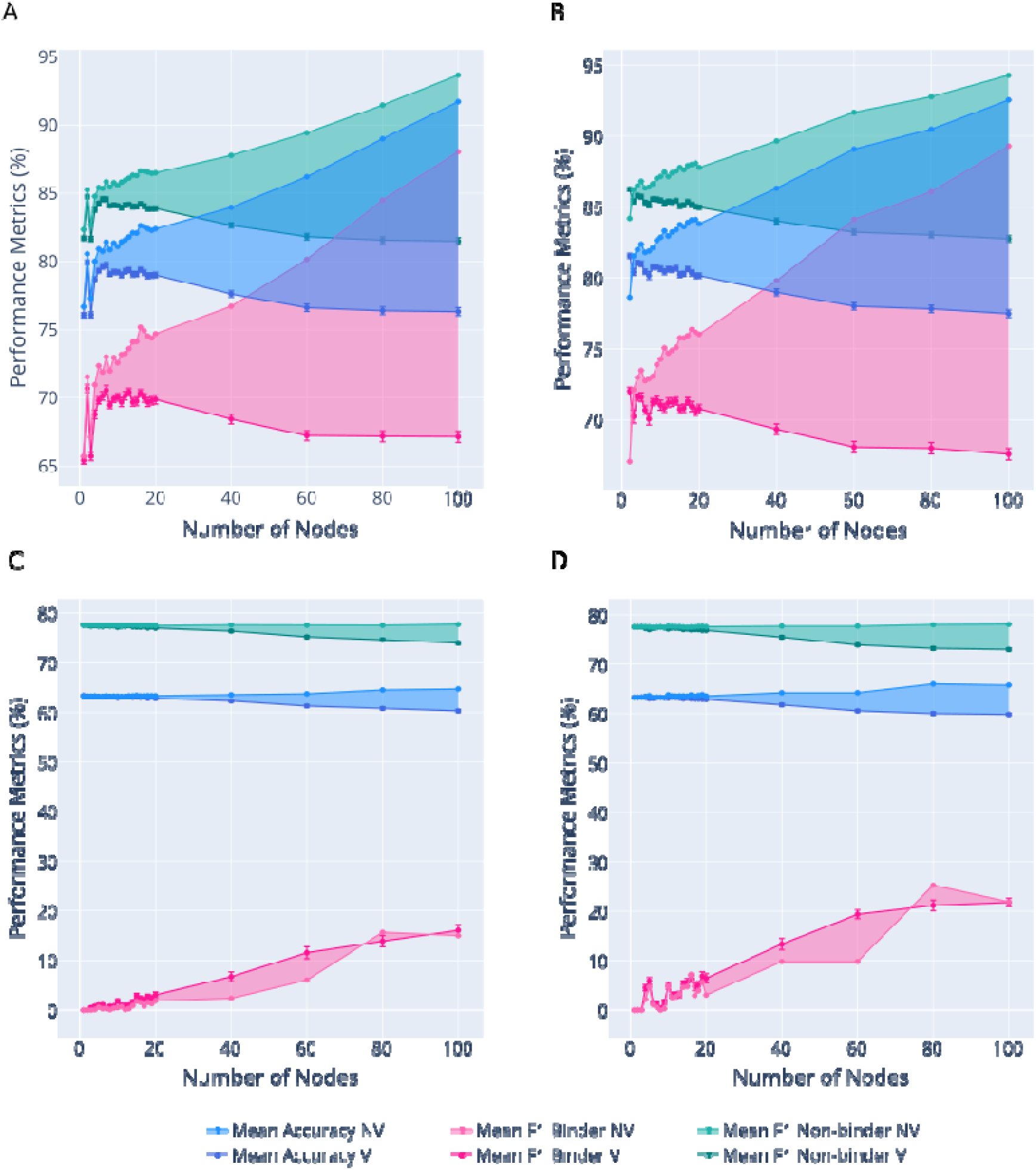
Performance metric value trends across a range of number of nodes (1 – 100) in different prediction models using: A. 15 atomic composition features (C-Coord2), B. 19 atomic composition features (C-Pos), C. Fourier Transform of non-uniform value representation of amino acids (FT-Seq), and D. Fourier Transform of uniform value representation of amino acids (FT-UniSeq). Performance metrics used are mean accuracy (blue), mean F1 values of ‘binder’ peptides (F1 Positive, in cyan), and mean F1 values of ‘non-binder’ peptides (F1 Negative, in pink). Filled areas between each performance metric show the difference between validated (V) and non-validated (NV) prediction results, with NV data points in lighter colours. All statistical values were calculated from 100 prediction runs; error bars represent the standard error of the mean.

Sequence-based methods evidently require a more complex network to enhance binder detection, as their F1 Positive score reaches a plateau around 0.20 (Figures 2C and 2D). The widening gap between the performance with and without cross-validation as model complexity increases suggests that the best-performing model may lack sufficient generalizability.

### Intuitive interpretation of the peptide preferences of survivin

The optimized network architecture (Figure 3) highlights two distinct pathways that lead to survivin binding decisions for a given peptide: Firstly, a pathway is created when many carboxyl groups and few amino groups are present which inhibit and activate the inhibitory perceptron 2, respectively. Secondly, amino groups activate the stimulating perceptron 1 and if simultaneously the number of carboxyl groups is low, survivin binding is likely predicted. Interestingly, phenyl rings is a supportive factor when paired with carboxyl groups as their presence inhibits perceptron 2 which in turn supports survivin binding decision. All atom types contribute to the prediction, although the weighted edges of features below 0.5 or exceed –0.5 are not displayed in Figure 3.

**Figure 3.**
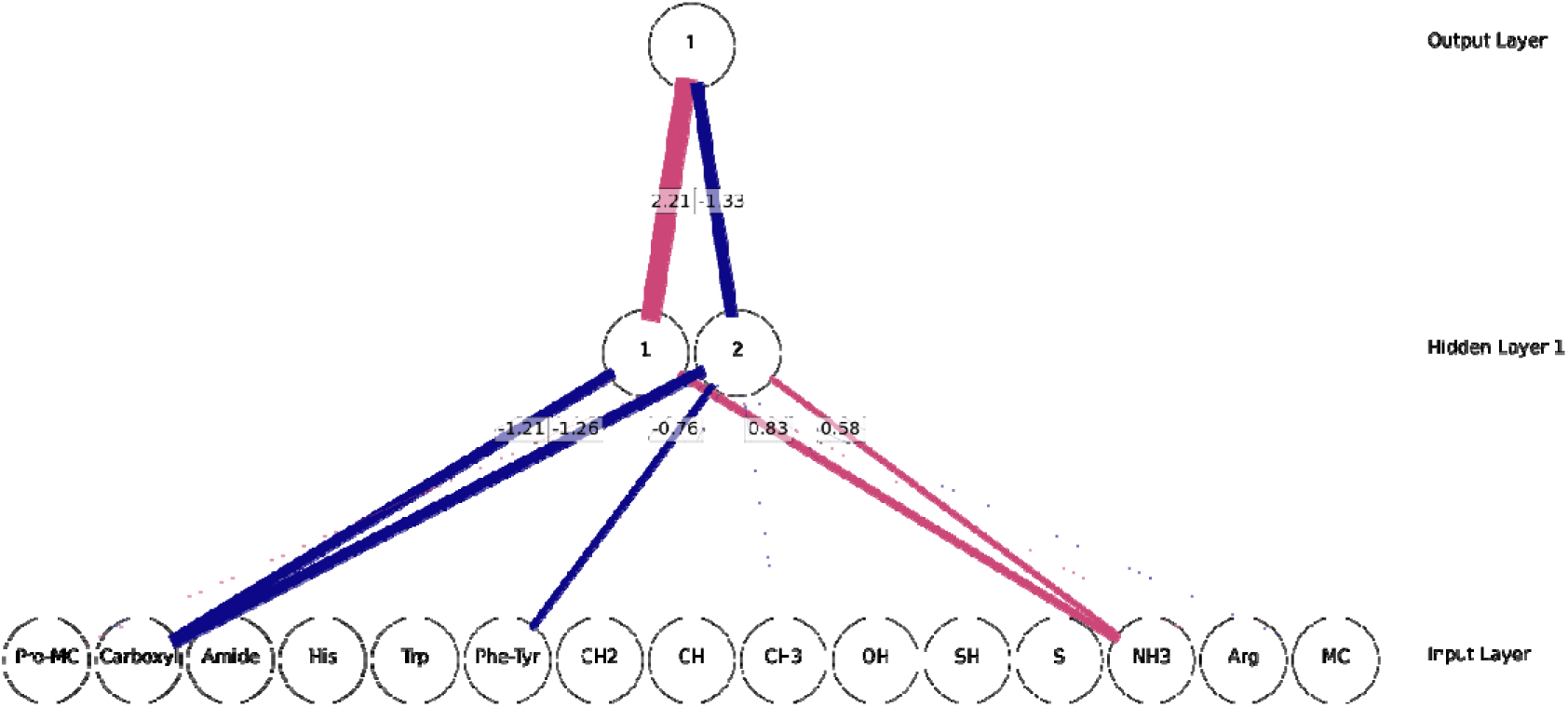
Multilayer perceptron architecture (C-Coord2 based) used in the prediction model for survivin-microarray peptide interactions. The input layer nodes correspond to various features (atom types). These features, listed from left to right, encompass distinct atoms of the proline main chain, carboxyl group, amide group, imidazole ring, indol ring, phenyl ring, methanetriyl-, methylene-, methyl-, hydroxyl-, sulfhydryl group, thioether of methionine, amino group of lysine, guanidino group, and standard main chain atoms. Positive weights are represented by pink lines, and negative weights by blue lines.

The occurrences of amino acids in peptide sequence are inherently anticorrelated since their numbers can be approximated well with a multinomial distribution. Each amino acid comprises various functional groups, some promote survivin binding while others inhibit it, allowing closely related amino acids, i.e. aspartate and glutamate, to be discerned based on their counts of methylene groups.

### The contribution of the peptide primary structure

To improve the sequence-based predictions, we employed bidirectional LSTM^45^ networks and embedding techniques. The most effective architecture had two LSTM layers, with 90 neurons in the first layer and 70 neurons in the second. The model comprised 243221 trainable parameters, demonstrated an accuracy of 85.7±0.1 % (cross-validated) and 89.7±0.2 % (non-validated).

Upon comparing the best composition-based and sequence-based predictions, we can infer that the composition alone cannot account for approximately 5% of the accuracy observed in the sequence-based method. However, we must still account for position-related artifacts. Overfitting was likely due to the low data to parameter ratio, as shown by the performance gap between the cross-validated and non-validated trainings. Given the Poisson counting errors in fluorescence detection, which can mislabel binders as non-binders, the high performance shown in non-validated training is unrealistic. Test set contamination^46^ due to peptide overlaps may also inflate the cross-validation performance, which can force the model to detect exact sequences instead of real physicochemical patterns. When using only every 3^rd^ peptide the accuracy drops to 79.2±0.2% (cross-validated) and 89.7±0.2% (non-validated).

The performance gap between cross-validated and non-validated sets, along with the diminished performance compared to the complete dataset, underscores the challenge of test set contamination and overfitting within the more intricate LSTM architecture. Mitigating contamination increased the network generalizability while reducing prediction accuracy to the level comparable with the composition-based prediction. The contrast between the composition-based and sequence-based networks is more pronounced when we consider the number of trained parameters for optimization, with 40 parameters (C-Pos) in composition-based network and 243221 in sequence-based LSTM model. Following Occam’s razor principle, the model which has 6000 times less parameter should be preferred when performance is similar. While larger datasets would enhance the LSTM model performance, obtaining extensive experimental biochemical data often remains a formidable challenge.

We also assessed the impact of amino acid position in the peptide sequence, as shown in Figure 4, where the weights are extracted as a function of position. Despite having only 35 parameters, the accuracy is 79.7±0.0% (cross-validated) and 80.1±0.2% (non-validated), comparable to composition-based methods. Figure 4A shows that tyrosines, aspartates, tryptophanes and arginines contribute the most to the prediction, in partial agreement with the composition-based model (Figure 3).

**Figure 4.**
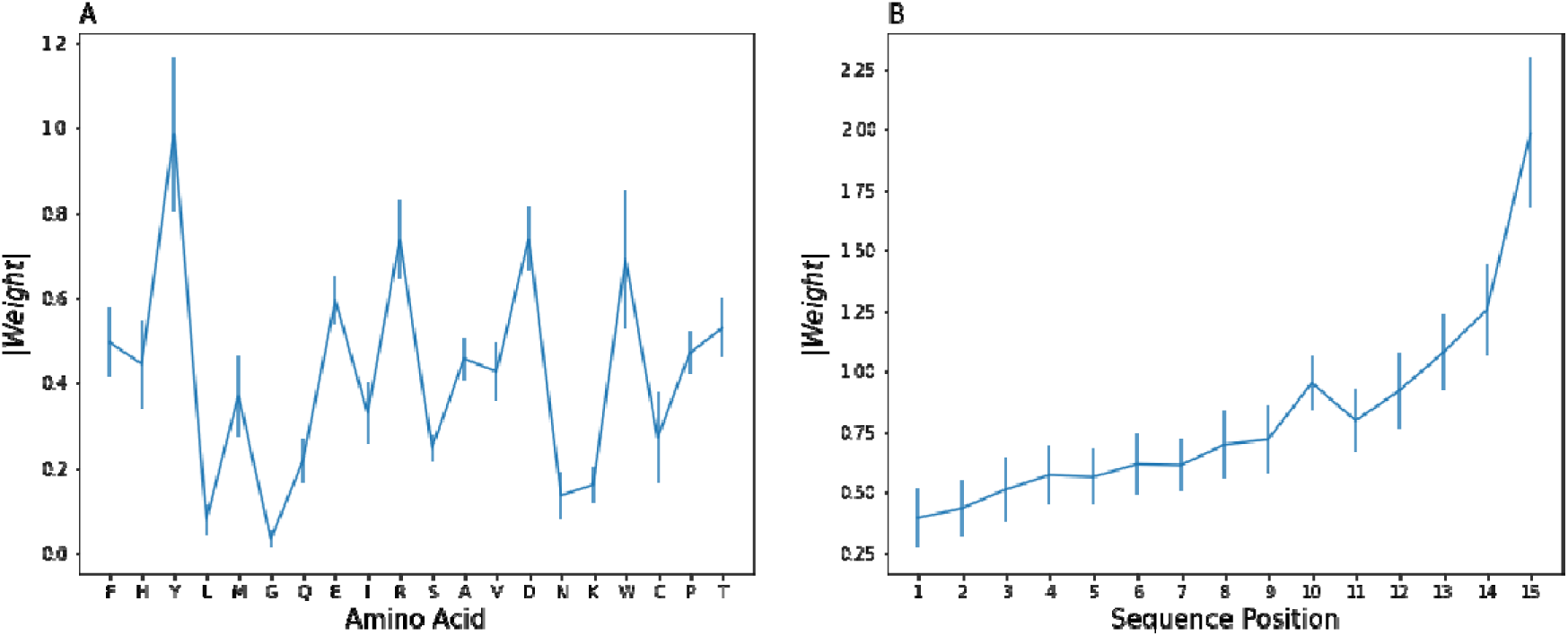
Position specific weights of the dense layer connecting the 2D embedding layer to the output. (A) Absolute value of the embedding quantity associate with every amino acid. (B) Absolute value of weights in the dense layer corresponding to the amino acid position from N to C terminus. The mean and the standard error of the mean are displayed based on 100 randomizations.

The gradual increase in weights from N- to C-terminus (Figure 4B) is likely linked to how the C-terminus of the peptide is tethered to the substrate through a linker, while the N-terminus extends freely into the solvent^47^. The relative importance of the C-terminus is unexpected because the surface exposed N-terminus intuitively has closer contact with survivin in solution. Such a position specific pattern is unlikely to represent a universal mechanism guiding survivin interactions in vivo, but these inherent experimental biases need to be addressed in the composition-based analysis.

### Visualizing the structure of peptide composition and sequence space and its connection to function

To gain a more intuitive insight to the advantage of the composition-based prediction, we visualized the peptide distribution according to the similarities between atom type composition using T-distributed stochastic neighbour embedding (t-SNE)^48^, a useful tool to map the multiple predictive features into a more intuitive 2-dimensional figure. Representing peptides based on their composition results in well-separated clusters (Figure S1A) with distinct separation between binders and non-binders, while only two clusters observed using the FT-Seq representation (Figure S1B) with binder and non-binder peptides mostly overlap with each other. Varying the perplexity value between 5.0 and 50.0 did not have a significant impact on the data point distribution^49^. While one might assume that peptides from a single protein share similar compositions, our clustering analysis challenges this perspective. Peptides from a single protein can be present in more than five recognizable clusters, exemplified by protein EZH2 in Figure S2, suggesting that a protein can contribute to multiple functions, as regions with different compositions are held together covalently in the polypeptide chain. We also discussed potential pitfalls of assay related errors in the accompanying text under ‘Statistical and experimental artifacts’, the quantitative connection between fluorescence intensity and atom type composition using t-SNE in Figure S3 and the accompanying text under ‘Quantitative binding representation and its relation to composition’.

### Analysis of a known survivin complex and structurally characterized interaction partners

Using the composition-based model, we investigated the impact of point mutations within the context of the surrounding residues by creating a metric M_bind_(n), where ‘n’ represents a residue position along the sequence based on the number of point mutations that result in peptides containing the given residue position being predicted to bind to survivin. M_bind_(n) of one indicates a region always predicted to bind survivin even if the position is mutated to any other amino acid, whereas zero indicates that no point mutation of the site can convert the region to a survivin binding region.

Through this approach, the wild-type amino acid is not given preferential treatment because it is assumed that the choice of a particular amino acid at a given position is often (not always) arbitrary, and alternative amino acids may perform equally well. This is because functionally neutral point mutations occur frequently during evolution^50^. In contrast, the composition-based context is robust, resistant to change, and evolves slowly over time. This is because it requires multiple point mutations with a clear drift direction before any substantial change in composition becomes apparent.

Figure 5 illustrates M_bind_(n) as a function of residue number, and it is evident that M_bind_(n) does not change smoothly along the sequence. Boundary between survivin binding and non-binding regions tend to be sharply defined and robust binding regions are focused to shorter segments. Discontinuous jumps are also prevalent. If an amino acid residue in the binding region has a significantly lower M_bind_(n) than its surroundings, it suggests that the wild-type residue at position n is crucial for maintaining the affinity of the surrounding region. This may lead to a loss of function (binding) with point mutations at that specific position. Conversely, if an amino acid residue in a non-binding region has a M_bind_(m) significantly higher than its surroundings, it indicates that the wild-type residue at position m is unsuitable for survivin binding within the given context. Position m may be associated with gain-of-function mutations resulting in survivin binding.

**Figure 5.**
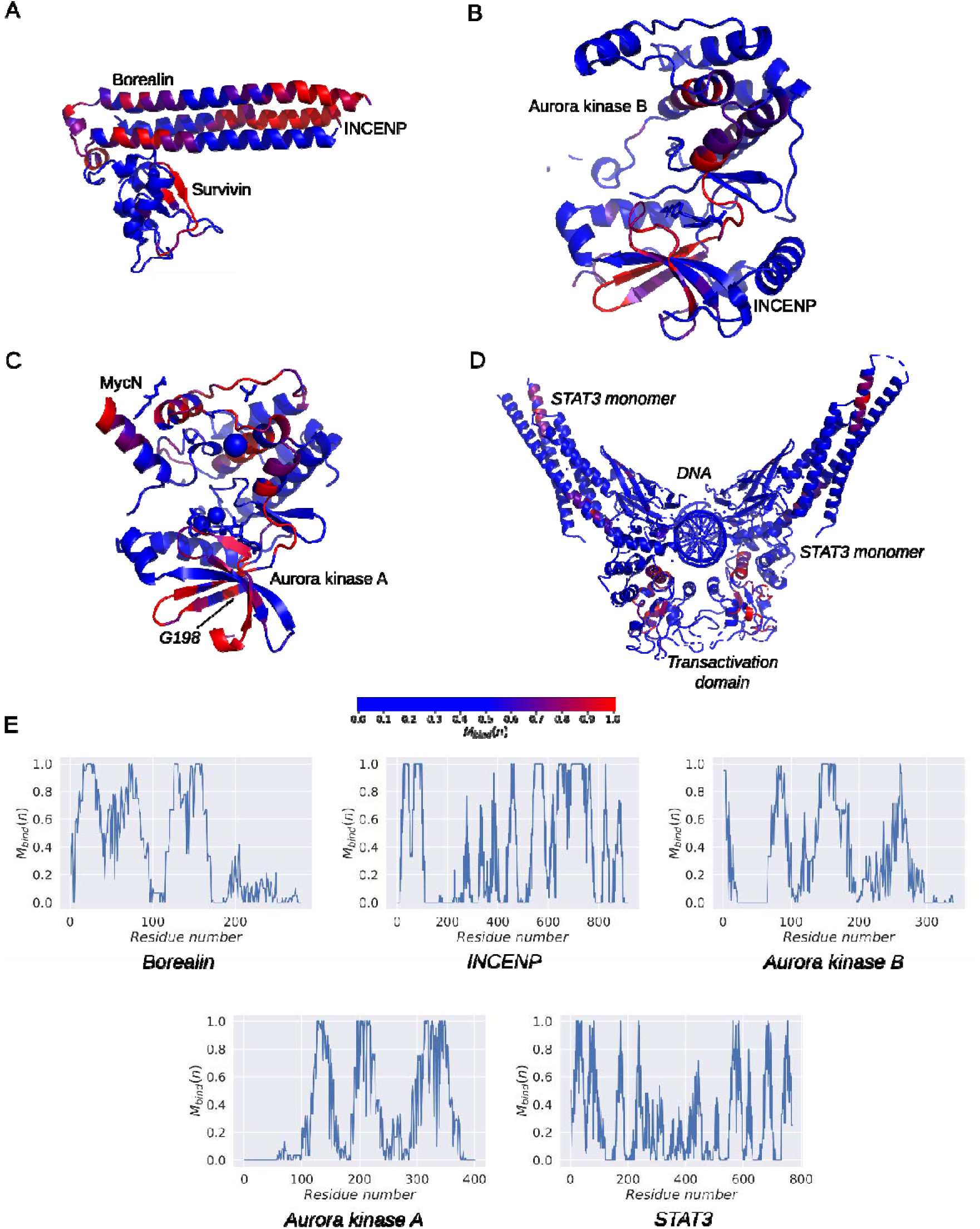
M_bind_(n) mapped on the ternary structure of known interaction partners of survivin: (A) CPC core consisting of borealin and INCENP (*PDB ID 6YIE ^95^*), (B) INCENP and Aurora kinase B (*PDB ID 4AF3 ^96^*), (C) Aurora kinase A and MycN (*PDB ID 7ZTL ^97^*), and (D) STAT3 bound to DNA (*PDB ID 6QHD ^60^*). The blue-red color map visualizes the M_bind_(n) range from 0.5 to 1.0. (E) The one-dimensional plot of M_bind_(n) the subunits INCENP, Borealin, Aurora kinase A, Aurora kinase B, and STAT3 is shown as a function of residues number.

Out of all identified survivin interaction partners, only the ternary complex of CPC involving survivin, borealin, and INCENP^51^ has been structurally characterized (Figure 5A). This complex is connected via N-terminal segments of borealin and INCENP, together with coiled-coil of survivin. While the helices of survivin and INCENP align in parallel, the helix of borelin lies anti-parallel to them. Survivin is predicted to bind to colocalized regions of borealin and INCENP (residues 27-46 in INCENP and 15-29 in borealin, in opposite directions). Borealin exhibits another predicted high-affinity region (59-76), which is observable in the crystal structure. Notably, INCENP and, to some extent, borealin, possess robust binding sequences compatible with survivin at other locations, which may come into short-range contact with survivin during their interactions.

Aurora kinase B plays an important role in phosphorylating CPC subunits, including survivin (Figure 5B). Survivin may also equally target Aurora kinase B active site, rendering it a favourable substrate. Aurora kinase B and A share 70% identity in the catalytic domain, yet the point mutation G198N transforms Aurora kinase A into B in terms of their specificity to interact with INCENP and survivin^52^. The high M_bind_(198) of Aurora kinase A (Figure S4), flanked by substantially lower M_bind_(197) and M_bind_(199) from tyrosine residues (ILRL**Y**<u>G</u>**Y**FHDA), suggests that replacing glycine with many other residues enhances survivin binding in that region, while removing even one tyrosine likely diminishes survivin affinity.

STAT3 is another survivin interacting partner^53^, which exhibits no extensive regions with high survivin binding, seeing almost no M_bind_(n) value reaching 1.0 (Figure 5D). However, both the N- and C-terminal regions generally show elevated M_bind_(n), with the latter encompass the transactivation (responsible for survivin interaction^53^) and SH2 domains near the DNA^54^, which is localized by M_bind_(n) with precision. The first helix of the coiled coil domain also displays heightened M_bind_(n)^54^.

### Proteome-wide survivin binding prediction and overview of survivin functions

We applied our survivin binding prediction model on the entire human proteome, leveraging the UniProt database. A higher binding ratio parameter and higher number of predicted binders was used as a parameter to judge a higher likelihood for affinity to survivin. The proteins with the higher than R_bind_ were used in enrichment analysis of biological annotations using Metascape^55^ and the network representation was mapped (Figure 6). Nuclear, cytosolic, and mitochondrial proteins, including members of the respiratory chain, are present among survivin binders, which is in accordance with the fact that survivin is firmly established to localize within these compartments. As survivin is anti-apoptotic, it is encouraging to see the group of enriched apoptotic proteins including caspase 6 and 8, Death-associated protein kinase 3 (DAPK3), AKT1/AKT3, Stratifin, IL18, High mobility group box 1 and 2 (HMGB1/2), Tyrosine 3-Monooxygenase/Tryptophan 5-Monooxygenase Activation Protein Zeta (YWHAZ), SEM1 26S Proteasome Subunit and Occludin (OCLN). These apoptotic proteins are non-homologous, catalyzing different reactions, and belonging to different compartments, yet their composition renders them attractive to survivin. The mitochondrial Smac/DIABLO protein, knowns for its proapoptotic activity, has an R_bind_ of 0.27 below the median R_bind_ of the proteome (0.31).

**Figure 6.**
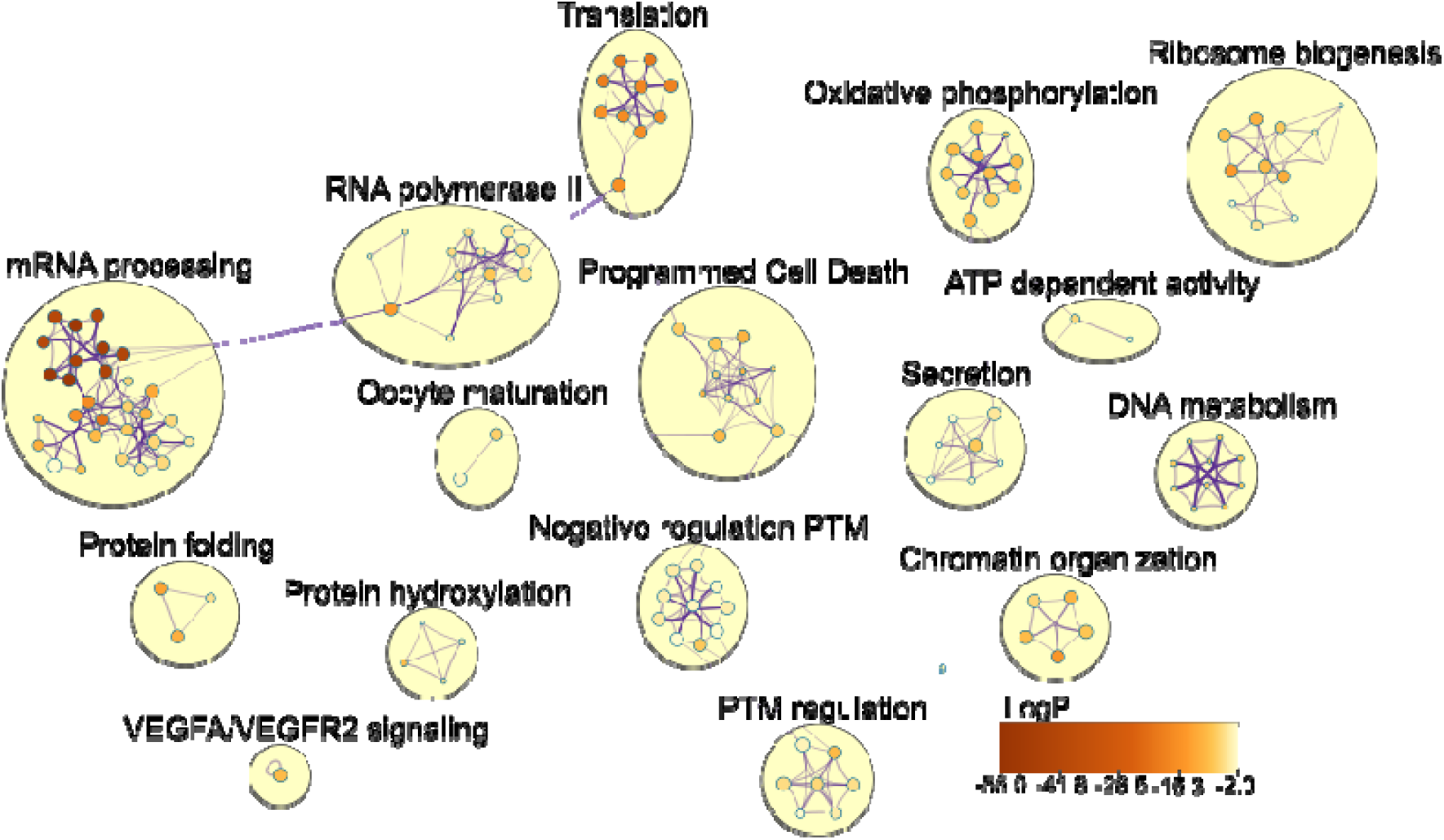
Annotation enrichment analysis conducted on human proteins using Metascape, focusing on proteins with an R_bind_ score exceeding 0.5, and excluding those with fewer than 5 constituting peptides (1396 proteins in total). The depicted network specifically highlights annotations with a p-value below 0.01 and an enrichment factor of at least 1.5.

Within the mitochondria, survivin influences oxidative phosphorylation^56,57^ and represented in Figure 6 by ATP synthase (subunits ε, ATP5MJ and K), NADH:ubiquinone oxidoreductase (B2, B5, B12, B14.5b, C2 and assembly factors NDUFAF1), succinate dehydrogenase complex (assembly factor 2), cytochrome b/c1 complex (subunits VI, VII, IX and X), cytochrome c oxidase (PET100, VIb2, VIc, VIIa1), ubiquinol-cytochrome-c reductase complex assembly factor 2.

Figure 6 also shows a significant number of translation-related proteins and proteins involved in RNA metabolism, transcription, and splicing, revealing putative functions of survivin. Recent experimental insights on survivin in micro-RNA biogenesis^58^ highlights the potential confirmation of binding predictions. Interaction of survivin with the cellular components are further discussed in the accompanying text under the heading "Additional observations concerning the Metascape analysis of predicted survivin binders of the proteome.”

A particular limitation of enhancement analysis stems from arbitrarily selecting the number of proteins enriched in survivin binding regions. Opting too few may overlook important survivin functions, while choosing too many may reduce the enrichment analysis sensitivity. Estimating the number of relevant survivin interaction partners is challenging with the information scarcity, although we can still potentially establish a lower limit. How survivin molecules, with its small solvent-accessible surface, engage in specific pairwise interactions also pose major question, which may be caused by our conceptual limitation by understanding PPIs as pairwise short-range interactions organized in pathways. The alternative in which survivin interacts with potentially all cellular proteins via long-range interactions is theoretically possible, but unattainable by statistical enrichment analysis. This radically different view on biological systems necessitates a discussion that probes whether there are physicochemical mechanisms in the background justifying the observed agreement. As we will discover, the answer is positive.

### Exploring the effectiveness of composition-based prediction

This study suggests exploring new approaches to understand and categorize interactions between proteins or biopolymers. While it is uncertain whether some fundamental concept may be missing, it is evident that variations in amino-acid compositions effectively capture structural and interacting protein features. Sequence and patterning information is certainly significant^59^, but it may not always offer a convenient representation for certain protein-related challenges. To explore new avenues, classical and quantum theories of collective excitations may be fruitful, as well as the physics of polymers and colloids with phase transition models. Current literature indicates the necessity for this cross-fertilization, as it is exemplified by the existence of intrinsically disordered proteins (IDP) or unstructured domains under physiological conditions^60^. Recent hypotheses propose that liquid-liquid phase separations might be inherent to proteins/polypeptides, irrespective of their sequence and structure (but with distinct interactions, electrostatic, hydrophobic, and H-bonding^61^). Understanding protein interactions at a compositional level can impact, vice versa, the study of phase transitions, such as the formation of nuclei via interactions with parent particles or existing nuclei in the bulk (secondary nucleation).

Initially, our study, departing from structural information, focused on atom displacement analysis in folded proteins. The observation that atoms with similar chemical properties exhibit shared displacement patterns^62,63^ led to a dynamic model in second-quantization, revealing coherent motion of phononic and dipolar collective excitations within the protein crystal^64^. This coherence extends between different proteins over intermolecular distances, involving identical functional groups. Testing this unconventional idea yielded a surprisingly simple and effective initial approximation^13^.

Motivated by the chemical principle "like dissolves like," our view extends the concept of miscibility between molecules, emphasizing atom counting and categorizing functional groups. The success of atom counting, particularly in predicting water miscibility, relies on identifying functional groups—collections of identical or different elements with a localized electronic structure. The C/O ratio, for instance, serves as a primary descriptor^65^ for inferring water solubility in organic molecules. However, exceptions to this trend exist^66^, and secondary selection criteria include molecular size and structure.

The miscibility concept does not lead to a clear binary choice between hydrophobic and hydrophilic compounds. This is illustrated by per- and poly-fluoroalkyl substances (PFAS), which don’t mix well with typical hydrophobic or hydrophilic substances. Our results suggest that the miscibility of simple molecules should parallel the interaction (or ‘affinity’) of proteins with different functional groups. Applying this logic to biology, protein interactions should be re-evaluated considering mechanisms leading to phase transitions or separations in condensed matter. Non-interacting proteins may not be neutral as well; their incompatibility can drive active structuring/segregation into separate phases. Analogies with polymer, colloid and interface sciences, such as "irreversible aggregation" or "reversible self-assembly" were proposed, with connections to diseases like amyloid fibrils formation^67^.

Despite several advancements, quantitative challenges persist. Understanding elementary mechanisms behind protein transitions remains tough due to compensating energy−entropy effects at various length scales^68^. The realistic free energy behaviour of macromolecular systems with heterogeneous composition is a non-local functional of monomer density distributions^69^, challenging the derivation of chemical potentials from phase diagrams. This complexity highlights the potential impact of more synoptic tools, such as the one based on atomic composition. However, the underlying explanation for empirical models may be more related to the dynamic behaviour of phases than their static properties alone. Common phase transitions can be associated with bifurcation theory, predicting significant perturbations in atomic dynamics with small alterations in control parameters^70^.

It has been suggested that proteins may operate near a critical point where distinct phases merge, given their exceptional functional sensitivity and mutational robustness^71^. This delicate balance would allow proteins to work at the edge of instability, demonstrating both high "plasticity" to small environmental fluctuations and high structural stability to maintain integrity. Formally, the critical temperature should depend on protein concentration, reaching a tricritical state when the folding temperature approaches the Q value^72^, akin to a continuous coil-globule transition in synthetic polymers^73^. Acknowledging features from phase transitions of both first-order (e.g. mass density) and second-order (entropy) seems to be essential in current theoretical interpretations. A first-order transition then can become continuous in the presence of a surface^74^.

A relevant example is the melting of a material, transitioning from an ordered solid to a liquid with disordered fluctuations. At the critical temperature, lattice vibrations significantly decrease, and the atomic/molecular constituents adopt a liquid-like diffusive regime. For a one-dimensional lattice of atoms, the equilibrium fluctuation of the mean squared displacement follows a spatial random walk^75^, akin to the law governing the end-to-end distance in a polymer as a function of molecular weight. This relation depends on factors like equilibrium spacing, temperature, and force constants connecting atomic pairs and discussed in the accompanying text under the heading ‘Fluctuations in a Monoatomic Lattice’. For a linear diatomic lattice with long-range oscillations (acoustic branch), the overall fluctuations turn out to be affected by the atomic composition by the mass proportions of their species, and this model is discussed in the accompanying text under the heading ‘Fluctuations in a Diatomic Lattice’. At the melting point, deviations from a perfectly periodic structure can be quantified while maintaining compositional-dependent long-range order. In reality, melting is a surface-initiated unstable process^76^, with a characteristic temperature approximately (20-30)% lower than the bulk value. Nevertheless, this remark does not invalidate the previous conclusions.

A representative mean field theory for phase separation in polymeric mixtures is Flory-Huggins’ (FH) and its variants like Overbeek-Voorn’s (OV) or Edmond-Ogston’s (EO). OV improves FH with long-range electrostatic interactions, relevant for complex coacervation ^77^, while EO is a (truncated) virial expansion^78^. FH assumes short-ranged, pointwise force fields without distinguishing between charged and neutral residues. OV does not consider the sign of charges. Random phase approximations (RPAs) may be used for a patterning description of charged units, provided the transition is not driven by strong critical fluctuations^79^. The key quantity remains the FH parameter (c), representing the energy cost for having lattice sites adjacent to the polymer occupied by solvent units, which we highlight in the accompanying text under the heading ‘Notes on Flory & Huggins’ (FH) solution theory’. Phase separation/aggregation occurs when the solvent is poor for the macromolecules (e.g. proteins rich in polar amino acids in water)^80^. When the segregation degree surpasses a critical value, the enthalpy contribution to separation prevails, resulting in a solution of (almost) pure phases in equilibrium. Discussing the magnitude of cN (N = polymer repeat unit number) aids in defining the character of segregation and the incompatibility unit at a microphase scale (e.g. the “oligo-nucleosomal clutch” in heterochromatin-like domains subjected to order-disorder transitions^81^).

In FH-like models, residues are treated as independent repeat units, lacking consideration for chain connectivity or molecular details, limiting the ability to address sequence- and structure-dependent interactions. Despite these constraints, this framework successfully balances energies contributing to phase separation^79,82^. Even a standard FH approach with average c accurately predicts the critical temperature for the germ granule protein Ddx4^83^. The phase transition rate is governed by the extent of concentration and conformational fluctuation, reaching the largest amplitudes at the critical point, inducing spontaneous demixing. Enhancing protein-protein interactions results in multiple energy minima, defining the composition of separated phases along the volume fraction coordinate^82^. In a mixture of neutral molecules in a macromolecular blend, the inflection point on the FH free energy curve determines a compositional-dependent critical point.

From the viewpoint of associative polymer physics, a biological molecule may be modelled as a sequence of stickers and spacers, carriers of attractive forces and non-attractive macromolecular segments, respectively^84^. This connection to associative polymer physics is discussed in the accompanying online text under the heading ‘Notes on Associative Polymer Physics and Percolation’. Stickers are expected to be short linear motifs (1–10 residues, SLiMs) in intrinsically disordered domains, or even single nucleotides in unfolded RNA molecules. This perspective arises from the idea that many membraneless biological condensates result from phase transformations and percolation phenomena^85^. Percolation occurs when, at a certain sticker concentration threshold, macromolecules form a system-spanning network. For a mixture of different polymers with stickers, the percolation threshold is composition-dependent, representing a measure of the attractive volume related to the sticker pair^84^.

Considering the number of identical atoms or functional groups as an order parameter reveals substantial changes when reaching a certain threshold in a given macromolecular domain. For instance in a peptide with 15 amino acids and a volume of 2000 Å^3^, having from 2 to 15 identical functional groups sets a concentration *c_eff_* ≈ *n_F_* / (
*N_F_ l_p_* 135 Å^3^) = (1.7 − 12.5) *M*, where *n_F_* is the number of functional groups or atoms and *l_p_* is the polypeptide length.^86^. This range, on the average, remains valid even though there is only one peptide molecule (i.e. an infinitely diluted solution). Adopting this approximation, if the number of functional groups is taken for simplicity as proportional to the polypeptide length, the effective concentration remains constant, regardless of the peptide or protein size. The concept of scale-freeness aligns with the use of atomic ratios as predictive tools, and a phase transition can occur in rather small protein segments of heavily biased compositions.

The behaviour of peptide or protein solutions at the local composition level differs significantly from low molecular weight liquids where all units are uniformly dispersed. Due to covalent bonding, residues are locally constrained to lattice-like structures influenced by force fields such as residue-residue and solvent-residue interactions (solvation, hydrophobic forces, weak interactions, etc.) as well as the amino-acid sequence. This implies that a peptide segment may exhibit a substantial compositional bias, potentially being as impactful as a similarly biased composition in a larger protein. However, when an equivalent number of identical amino acids are uniformly dispersed within a larger protein, it might not necessarily induce a phase transition. In compounds highly soluble and freely diffusing in a given environment, the solubility limit often falls within the former range (1.7-12.5 M), leading to precipitation and the emergence of highly ordered coupled motions reminiscent of those detected in soft matter and materials^87^. This process, when associated with a bifurcation point, could facilitate the folding of these structures into a 3D configuration, akin to the principles governing crystal nucleation & growth, highlighting a dynamic attractor resistant to perturbations. Alternatively, it may give rise to distinct periodic coordinated dynamics rather than incoherent motions. Dynamic transitions can also be initiated by control parameters like temperature changes or the application of a pulling force, as observed in unfolding^88^ or protein unfolding through pulling^39^.

Over billions of years, organisms have leveraged fundamental physical principles, exemplified by signal peptides like nuclear localization signals with significant compositional biases^89^. Such peptides, despite low information content, hold substantial abstract signal value due to their unique enrichment in specific functional groups, as seen in the PKKKRKV segment found in the SV40 Large T-antigen. Amino acid distributions, primarily determined by genetic material, are further influenced by covalent post-translational modifications like phosphorylation, acetylation, methylation, ubiquitination, nitrosylation, hydroxylation, sulfation and deamidation. These modifications, observed individually or in patterns like hyperphosphorylation, enable protein composition to adapt, triggering new collective dynamics based on introduced groups and the organism’s physiological state.

In summary, this work has focused on the predictive influence of elemental composition in protein-protein interaction systems, establishing its conceptual alignment with biological processes and molecular functions. Although finer mechanistic details await further elucidation and formalization, our identification of key questions signals the need for exploration to unravel the intricate organization of biological organisms.

## METHODS

### Peptide microarray experiments

The peptide microarray experiment was described previously^13^ and corresponding data is the basis of this analysis^90^. Briefly, peptide microarray was designed with a total of 36 proteins using PEPperCHIP Peptide Microarrays (PEPperPRINT Gmbh). A complete list of proteins and more detailed description of the method can be found in previous research^13^. Each protein sequence was divided into 15 amino acid peptide units, with 10 amino acid overlap. Background interactions was examined by pre-staining one microarray with the secondary 6X His Tag Antibody DyLight680 antibody (1:1000) and monoclonal anti-HA (12CA5)-DyLight800 control antibody (1:1000). Another peptide microarray was incubated with survivin at a concentration of 1 μg/ml and stained with the secondary 6X His Tag Antibody DyLight680 antibody (Rockland Immunochemicals, Pottstown, PA, USA) and the monoclonal anti-HA (12CA5)-DyLight800 control antibody (Rockland Immunochemicals, Pottstown, PA, USA). The read-out was performed using LI-COR Odyssey Imaging System with scanning intensities of 7/7 (red/green). HA and His-tag peptides were also stained simultaneously in the assay as internal quality control. PepSlide Analyzer was used for quantification of spot intensities and peptide annotation. The resulting data was stored as a table with information on protein identifier, peptide sequence, and fluorescence intensity.

### Development of feature scoring set for the representation of peptides

The features considered encompassed the standard four non-hydrogen main chain atoms (MC), except glycine (3 non-hydrogen atoms) and proline (2 non-hydrogen atoms). Additionally, specific features included the glycine Cα atom (CA-Gly), recognizing its methylene group nature instead of a methanetriyl-group. Proline-specific main chain atoms (Pro-MC) were considered due to the circular proline side chain linking the amide nitrogen to Cα, resulting in distinctive dynamics. The content of side chains were deconstructed to carboxyl groups of aspartate and glutamate (carboxyl), the amide groups of asparagine and glutamine (amide), the imidazole ring of histidine (His), the indole ring of tryptophan (Trp), the phenyl group of phenylalanine and tyrosine (Phe-Tyr), the phenolic hydroxyl group of tyrosine (OH-Tyr), the hydroxyl group of serine and threonine amino acids (OH), the sulfhydryl group or oxidized variants of cysteine (SH), the thioether of methionine (S), the amino group of lysine (NH3), and the guanidino group of arginine (Arg). Carbon atoms not part of the main chain, carboxyl, amide and aromatic functional groups were assigned to methyl- (-CH3), methylene- (-CH2-), and methanetriyl- (>CH-) groups. These categories resulted in 17 features, each corresponding to the number of non-hydrogen atoms included. (Table S1)

We also embarked on a stepwise simplification of the C-Coord1 model. Initially, we consolidated the features of phenolic and alkyl hydroxyl groups into a single feature. Additionally, we described glycine as comprising three standard main chain atoms and a methylene group, thus eliminating the necessity for a separate atom type for the Cα atom of glycine. This refinement resulted in the C-Coord2 model (Table S2), which incorporates 15 features. We further adjusted C-Coord2 model by restricting oxygen atoms in carboxyl and carbonyl side chains to be indistinguishable. Nitrogen atoms in non-heterocyclic side chains were compelled to be identical, and carbon atoms with no bonded hydrogens (C0) in amide, carboxyl, and guanidino side chain groups were also forced to be identical. These modifications led to the development of the C-Coord3 model, which encompasses 13 features. (Table S3)

Another method used here to characterize side chain carbon atoms involved considering their distance from the main chain (C-Pos, Table S4). The rationale is exemplified here by the varying dynamics of methylene side chain groups. For example, the Cβ atom of lysine forms a quasi-rigid group with the Cα, the main chain amide nitrogen, and carbonyl carbon, which in this (β) configuration mirrors the fluctuations of the main chain. On the other hand, the Cδ atom has a high degree of freedom, and its position is influenced by multiple χ torsion angles in the side chain. Suggesting that these two methylene groups exhibit similar behavior in models C-Coord1-3 could be an overly simplified abstraction when taking the dynamics of these atoms into account. The drawback of the C-Pos model is that number of features is 19 only one less than the number of naturally occurring amino acids and it does not distinguish between methyl- (-CH3), methylene- (-CH2-), and methanetriyl- (>CH-) groups. For comparison, the AA-Letter model (Table S5) was included to represent amino acid composition, assuming each amino acid is unique and possesses equal weight, resulting in a model with 20 features. We compared these four models with elemental composition models, where the features included the counts of hydrogen, carbon, nitrogen, oxygen, and sulfur atoms (5 features in “Chemical Formula”, Table S6).

For the characterization of the electrostatic and hydrophobic properties of peptides, we used two different approaches. In the Pos/Neg/HyPho model (Table S7), we approximated these properties by considering the number of basic and acidic amino acids and whether they were classified as hydrophobic, resulting in 3 features. In the NetChrg/SumHyPho model (Table S8), we determined the net charge by subtracting the count of acidic amino acids from basic ones and calculated the sum of hydrophobicity according to the Kyte/Doolittle scale^44^, yielding two features.

### Applying the feature representation to peptide microarray dataset

Different sets of feature scorings were developed based on atom type composition and applied on each amino acid in the dataset. Briefly, a feature scoring table was created by listing 20 amino acids, with each row representing each distinct amino acid.

For feature representation based on atom type composition, each amino acid is represented by a set number of scores, according to the number of categories present in the scoring set, as shown in Figure 1. The scoring set was then applied to each peptide sequence from the microarray dataset, summing the scores of each atom type category from all amino acids in a particular peptide.

For feature representation based on AAindex numerical values, each amino acid is represented by one value based on RACS820104 index^91^ from AAindex database^15^. Fourier transform was applied on this one-dimensional array of peptide representation using Fast Fourier Transform from scipy 1.6.2 package and only the real part of the Fourier Transform was considered.

### t-SNE dimensionality reduction analysis of peptide microarray dataset

To calculate the spatial distance between peptides, t-SNE module from scikit-learn 1.0.1 package was applied to both atom type composition and AAindex representations. In the latter case, t-SNE was performed on the representation before (Seq, as in Figure S1C and S1D) and after (FT-Seq, as in Figure S1B) Fast Fourier Transform step. The following parameters were used: n_components=2, perplexity=30.0, verbose=1, and random_state=123.

### Prediction of survivin binding based on the peptide microarray dataset

Peptides from the microarray dataset with non-zero fluorescence intensities were marked as binders, and the ones with zero intensities were marked as non-binders. 5388 peptides from the dataset were divided into equally large training and test sets. The model was built using multi-layer perceptron classifier from scikit-learn 1.0.1 on both sets with the following settings: solver=’adam’, learning_rate=’constant’, random_state=1, max_iter=10000, and activation=’relu’. Hyperparameter tuning was done on solver, learning_rate, max_iter, hidden_layer_sizes, activation parameters. Fine tuning of the number of nodes is shown in Figure 2. Prediction statistics were obtained from the averages of 100 prediction runs.

### Improving the sequence-based prediction of survivin binding using bidirectional LSTM

In bidirectional LSTM refinement, we made no alterations on the configuration similar to the one used for optimizing the dense layer, except by the addition or omission of one dense layer after the final bidirectional LSTM layer. We tested either the Keras Embedding Layer^92^ or ProtT5 encoding^93^, as well as testing up to two dense layers and two bidirectional LSTM layers. Additionally, the model incorporated dropout between layers to aid regularization.

To assess the impact of position, we utilized a straightforward model consisting of a one-dimensional embedding layer connected to a dense layer featuring a single intermediate layer neuron. This setup approximates a position-specific summation function. In contrast, our AA-Letter model has explicit summation of amino acids ignoring their positions.

### Highlighting predicted survivin binding regions and measuring the impact of point mutation in survivin binding proteins

The metric for point mutation impact on the surrounding residues M_bind_(n), where ‘n’ represents a residue position along the 15 amino acid peptide sequence, is based on the number of point mutations required to make said residue being predicted to bind to survivin. Due to 10-residue overlap between 15 residue-long peptides, a residue position is typically found in three unique peptides, and with 20 natural amino acid variants, we ended up with 60 peptide permutations, each with a binding prediction. The total number of mutations predicted to be binders divided by the total number of peptides with variants at ‘n’ equals M_bind_(n). The C-Pos model with two neurons in the intermediate layer was applied on this dataset and 90% of the microarray data set was used. The peptides with C-terminal cysteins were excluded from the data set. The tested peptides were scaled using the microarray data distribution parameters.

### Application to human proteome and gene ontology enrichment in predicted survivin binder proteins

Human proteome database was obtained from UNIPROT. The sequences of all proteins in the database were used to generate a dataset containing 15mer peptide units with 10 amino acid overlaps, mimicking the peptide microarray experiments. The tested peptides were scaled using the microarray data distribution parameters. The trained C-Pos model (90% of the data with C-terminal cysteines excluded) with two neurons in the intermediate layer was applied on this dataset and the results were consolidated to get the binding ratio of each protein (R_bind_, the number of binder peptides divided by the total number of peptides each protein contains). The list of human proteins and their associated R_bind_ value is listed in Table S10. The highly ranked proteins were subjected to biological annotation enrichment analysis using Metascape (https://metascape.org)^55^. Network mapping of enrichment annotation terms from predicted survivin interaction partners was made using the Cytoscape platform^94^.

## Acknowledgements

This work has been funded by grants from the Röntgen-Ångström Cluster Framework of the Swedish Research Council (G.K., 2015-06099), the Swedish Research Council (M.I.B., 2017-03025), the Swedish Association against Rheumatism (M.I.B., R-566961, R-751351 and R-860371), the King Gustaf V:s 80-year Foundation (M.I.B.), the Regional agreement on medical training and clinical research between the Western Götaland county council and the University of Gothenburg (M.I.B., ALFGBG-717681 and ALFGBG-965623). This project has received funding from the European Union’s Horizon 2020 research and innovation program under grant agreement No 964203 (Long-range electrodynamic INteractions between proteinS — LINkS). S.A.M. acknowledges financial support from HrZZ (Croatian Science Foundations, Grant ZFM – 0352Z32 – HZZ – POTENTIALS – 3456).

## Author Contributions

G.K. and S.A.M. conceptualized the study outline and discussion. G.K. supervised the research. A.L.A., T.N.O., M.J. and M.J.-G.B. and G.K. analyzed the results from the peptide microarray. M.I.B. and S.P.W. contributed to the discussion regarding the cellular function of survivin. The manuscript was prepared by A.L.A., T.N.O., S.A.M., and G.K. with additional input from all authors.

## Declaration of interest

G.K. and M.I.B. submitted a patent application for the machine learning method described in the paper. The remaining authors have no competing interests.

## Inclusion and Diversity

We support inclusive, diverse, and equitable conduct of research.

## Notes

### Summary of Updates

Revision was made on the following documents: - Main manuscript (revised texts in the Results and Discussion section to minimize potential ambiguities, revised Figure 1, Figure 2, Figure 5, Figure 6, and Table 1 as well as adjusting the text according to the revised Figures and Tables) - Supplementary material file (corrected the texts referring to Figures and Tables in the Supplementary material and removed Figure S5) - Competing interest (corrected the statement as 2 authors submitted a patent application for the machine learning method described in the paper

