## Supplementary material for "Deciphering Peptide-Protein Interactions via Composition-Based Prediction: A Case Study with Survivin/BIRC5": Table S1, Table S2, Table S3, Table S4, Table S5, Table S6, Table S7, Table S8, Table S9, Figure S1, Figure S2, Figure S3, Figure S4

*Statistical and experimental artifacts*

Another issue of using numerical value representation of amino acid sequence (Seq and FT-Seq) is the scale used for representing amino acids. Figure S1C shows the peptide distribution when the spacing between the numbers separating adjacent amino acids are not uniform. Here, the 20 amino acids are represented by numerical values between 0.41-3.14, with 18 amino acids having rather uniformly separated values between 0.85-1.75. One may already note the extreme values given to methionine (0.41) and cysteine (3.14). Looking at Figure S2C, we found that one ‘small island’, consists of only binding peptides, comes from peptides with cysteine in position 15 in the sequence. Each of the other ‘small islands’ also belongs to each position along the 15-amino acid peptide sequence in our dataset with just one cysteine present at that position. Peptides outside the small islands and the ‘main continent’ contain more than one cysteine, some of which may be at the C-terminus. The clustering therefore may not be biologically relevant, unless the presence of a single cysteine at a specific location is crucial for a highly specialized niche function. FT-Seq and mapping with a uniform spacing of value representation (UniSeq) eliminate the possibility to detect this propensity. The numerical representation represents fractional occurrence of amino acids within structures characterised by a left-handed turn in four-Cα scale and located in the extended regions of proteins. Rackovsky and Scheraga identified six distinct polypeptide backbone structures of four-Cα segments according to curvature (κ) and torsion (τ) angles^1-3^ to visualize longer-range interactions beyond single residues^1,2^. Applying appropriate fractional occurrence values to residues according to their context in their four-Cα unit backbone structures may lead to a better prediction.

We learned an unexpected lesson from Figure S1C that classification can depend on trivial factors like the presence of 15 islands for cysteine at a specific position, which can be modelled with a simple decision tree instead of a complex architecture. We also found an assay-specific artifact: C-terminal cysteines, despite being the farthest from the contact area, enhance survivin binding four times more than N-terminal residue (Figure 4B), but C-terminal cysteines rarely present *in vivo*. Model prediction using the reduced dataset (5287 peptides instead of 5385), eliminating peptides with C-terminal cysteines, yields 80.6% mean accuracy, 71.0% mean F1 of binders, and 85.5% F1 scores of non-binders, comparable to the full dataset (Table 1). Simple neural networks designed with physical and chemical intuition are less prone to functionally irrelevant bias compared to more complex ‘blackbox’ networks.

*Quantitative binding representation and its relation to composition*

We predominantly discussed classification performance. We tested regression models, but we were not successful achieving convincing correlation between the predicted and observed fluorescence intensities. Nevertheless, we visualized the connection between the fluorescence intensity and the atom type composition using t-SNE. Figure S3A shows regions where strongly binding peptides are accumulated, highlighted inside coloured rectangles.

Next, we wanted to investigate whether only few features used in the t-SNE distribution map are significant to determine peptide binding to survivin, which should be shown by an overlap of the number of each feature with binding intensity. The number of atoms belonging to individual features do not correlate strongly when mapped t-SNE representation. Atoms belonging to carboxyl groups are somewhat exceptional and highlighted in Figure S3B. The rectangles in Figure S3B indicate the peptides enriched with carboxyl groups which coincide with high fluorescence regions in Figure S3A. In some cases, the highlighted regions with high fluorescence intensity on the lower part of Figure S3A have below average content of carboxyl groups. Although carboxyl groups can be important for survivin, but they are not the only deterministic feature to infer peptide binding to survivin.

*Additional observations concerning the Metascape analysis of predicted survivin binders of the proteome.*

Consistent with previous experimental findings^4,5^, predicted survivin binding is also enhanced in clathrin components, including Clathrin light chain A and B (Lca, Lcb) and several adaptor protein 1 components. Chromobox (CBX) family proteins^6^ are also highly ranked on the R_bind_ scale which exhibit an affinity for methylated lysines in histone N-termini, making some CBX proteins closely linked to PRC1 which recognizes H3K9me3 signals. They are crucial for embryonic development and regulating heterochromatin formation, by repressing the expression of genes associated with differentiation, thus contributing to stem cell identity maintenance^6^. Survivin has also been recently recognized as a negative regulator of PRC2 by associating with EZH2 subunit^7^. CBX3 (Q13185), CBX5 (P45973), CBX1 (P83916) have high R_bind_. However, CBX2, CBX4, CBX6, and CBX8 are known to associate with PRC1^8^ but do not have high R_bind_. CBX1, CBX3 and CBX5 have a distinct function in their interaction with lamin B receptor (LBR)^9^, and CBX7 is the only one that is associated with PRC1^10^. Survivin appear to have a high-precision sensitivity to distinguish the composition of different homologous CBX proteins, emphasizing that composition-based screening is adequate for discerning closely related cellular functions. The method can identify which proteins within a family are involved in different complex types.

Troponin subunits and tropomyosins are also predicted to bind survivin indicating a hitherto underappreciated link to the actin cytoskeleton. Tropomyosin forms a coiled coil and survivin is also known to engage in coiled coil formation together with INCENP and borealin. Tropomyosin-1 (Q8NBA8), -2 (P0DKB5), -3/-5 (Q12816), and -4 (Q13641) all belong to the top 2% survivin binders, indicating an underexplored link to the actin cytoskeleton, cancer metastatis, and several autoimmune diseases^11-13^. The decreased expression of HMW tropomyosin-2/-5 isoforms and increased expression of LMW isoforms are linked to metastatic phenotype^14^. Survivin forms a coiled coil with INCENP and borealin, a feature also found in tropomyosin, besides having rich charged residues, which is sufficient to robustly provoke an autoimmune response and favourable for survivin binding^15^.

*Fluctuations in a Monoatomic Lattice*

As a simplest prototype model, consider a monoatomic lattice of $N$ unit cells for $N$ atoms in one dimension, each taking a position $\boldsymbol{n}=n\boldsymbol{l}$**,** with $n=1, \ldots N$ and spacing $l=|\boldsymbol{l}|$ (unit cell size). We represent the displacement of the $n$-th atom from its equilibrium position in Fourier’s series of the wave vectors in Brillouin’s reciprocal space, obeying $-\pi\leq\boldsymbol{k}\cdot\boldsymbol{l}<\pi$:

$\xi_{\boldsymbol{n}}=\frac{1}{\sqrt{N}}\sum_{\boldsymbol{k}} q_{\boldsymbol{k}}e^{i\boldsymbol{k}\cdot\boldsymbol{n}}$	(S1)

where $q_{\boldsymbol{k}}$ are the Fourier components. Its squared displacement, from any atom chosen as scratch ($\boldsymbol{n}=\boldsymbol{0}$), thus writes as:

$\left( \xi_{\boldsymbol{n}}- \xi_{\boldsymbol{0}} \right)^{2}=\frac{1}{N}\left[ \sum_{\boldsymbol{k}} q_{\boldsymbol{k}}\left( e^{i\boldsymbol{k}\cdot\boldsymbol{n}}\boldsymbol{-}1 \right) \right]^{2}$	(S2)

This sum runs evidently within $-\frac{\pi}{l}\leq k=|\boldsymbol{k|}<\pi/l$, the spacing in the one-dimensional reciprocal lattice being inversely proportional to the total length of the atomic system, $|\Delta\boldsymbol{k}|=2 \pi/L$, with $L=Nl$. To thermally average Eq. (S2), we recall the equipartition theorem, applied to the potential energy per mode $\boldsymbol{k}$ of a one-dimensional crystal with atomic mass $m$:

$\left\langle u_{\boldsymbol{k}} \right\rangle=\frac{m}{2}\left\langle q_{-\boldsymbol{k}}q_{\boldsymbol{k}} \right\rangle\Omega_{\boldsymbol{k}}^{2} =\frac{k_{B}T}{2}$	(S3)

Accordingly, for any couple of Fourier’s components:

$\left\langle q_{\boldsymbol{k}}q_{\boldsymbol{k}'} \right\rangle= \frac{k_{B}T}{m\Omega_{\boldsymbol{k}}\Omega_{\boldsymbol{k}\boldsymbol{'}}}\delta_{\boldsymbol{k},-\boldsymbol{k}'}$	(S4)

where $\delta$ clearly is Kronecker’s symbol and $\Omega_{\boldsymbol{k}}= \Omega_{-\boldsymbol{k}}$. Applying Eq. (S4) into the squared expression returns, upon the property ${(\sum_{k} a_{k})}^{2}= \sum_{ij} a_{i}a_{j}$:

$\left\langle\left( \xi_{\boldsymbol{n}}- \xi_{\boldsymbol{0}} \right)^{2} \right\rangle\equiv{\delta R}_{\boldsymbol{n}}^{2}=\frac{{2k}_{B}T}{M}\sum_{\boldsymbol{k}} \frac{1-\cos(\boldsymbol{k}\cdot\boldsymbol{n})}{\Omega_{\boldsymbol{k}}^{2}}$	(S5)

### $M=Nm$ being the total mass. The sum over reciprocal vectors can now be replaced by a linear integral in the continuum space, with density $1/|\Delta\boldsymbol{k}|$:

${\delta R}_{\boldsymbol{n}}^{2}\approx\frac{k_{B}Tl}{\pi m}\int_{-\frac{\pi}{l}}^{\frac{\pi}{l}} \frac{1-\cos(knl)}{\Omega_{\boldsymbol{k}}^{2}}dk$	(S6)

Peierls’ result relies^16^ on i.) approximating the cosine function to second order in $k$, ii.) working in the long-wavelength limit, $\Omega_{\boldsymbol{k}}\approx lk\sqrt{\gamma/m}$ and, especially, iii.) noting that the integral in Eq. (S6) gives a dominant contribution in $-1/(nl) <k \leq$ $1/(nl)$. After such substitutions, a random walk law:

${\delta R}_{\boldsymbol{n}}^{2} \approx na^{2}$	(S7)

turns out with step length $a=\sqrt{k_{B}T/\pi\gamma}$. Here, $\gamma$ accounts for the (harmonic) spring constant between adjacent units, and depends on the atom composition, while it is intriguing to note that $l$ does not take part in Eq. (S7).

The trend of $\delta R_{n}^{2}$, quantifying the deviation from a perfectly periodic structure, is expected to hold at the melting point as well, providing further indications for the existence of a compositional-dependent long-range order in a phase transition.

*Fluctuations in a Diatomic Lattice*

In a diatomic lattice^17^, two atomic species with masses $m_{i}$ ($i=1,2$) are to be dealt with. In this case, it is convenient to start from:

$\xi_{\boldsymbol{n}i}=\frac{1}{\sqrt{Nm_{i}}}\sum_{j,\boldsymbol{k}} e_{i}^{j}\left( \boldsymbol{k} \right)Q_{\boldsymbol{k}j}e^{i\boldsymbol{k}\cdot\boldsymbol{n}}$	(S8)

### summing the new Fourier’s components ($Q$) over the vibration branches ($j$). The real terms $e_{i}^{j}$ characterize the displacement of the mass $i$ in the branch $j$, are even functions of $\boldsymbol{k}$, and obey:

$\sum_{i} e_{i}^{j}\left( \boldsymbol{k} \right)e_{i}^{j'}\left( \boldsymbol{k} \right)= \delta_{j,j'}$	(S9)

### With unlike atoms in the lattice, the potential energy gets generally dependent on a force matrix, $F_{ii'}\left( \boldsymbol{k} \right)$, with the properties $F_{ii'}\left( \boldsymbol{k} \right)= {F_{i'i}}^{*}\left( \boldsymbol{k} \right)= {F_{ii'}}^{*}\left( \boldsymbol{-k} \right)$ ($i\neq i'$) and:

$\sum_{i'} F_{ii'}\left( \boldsymbol{k} \right)e_{i'}^{j}\left( \boldsymbol{k} \right)= \Omega_{\boldsymbol{k}j}^{2}e_{i}^{j}\left( \boldsymbol{k} \right)$	(S10)

where $\Omega_{\boldsymbol{k}j}$ yields the dispersion law specific to the branch $j$. As a consequence of Eq. (S10), let $n_{c}$ be the number of atoms per unit cell, the equipartition law in Eq. (S3) becomes:

$\left\langle U_{\boldsymbol{k}} \right\rangle=\frac{1}{2}\sum_{ii^{'},jj^{'}} F_{ii'}e_{i}^{j}\left( \boldsymbol{k} \right)e_{i'}^{j'}\left( \boldsymbol{k} \right) \left\langle Q_{-\boldsymbol{k}j'}Q_{\boldsymbol{k}j} \right\rangle=\frac{{n_{c}k}_{B}T}{2}$	(S11)

and, on exploiting Eq. (S9) and Eq. (S10), Eq. (S4) transforms into:

$\left\langle Q_{\boldsymbol{k}\boldsymbol{,}j}Q_{\boldsymbol{k}'\boldsymbol{,}j'} \right\rangle= \frac{n_{c}k_{B}T}{\Omega_{\boldsymbol{k}j}\Omega_{\boldsymbol{k}\boldsymbol{'}j'}}\delta_{\boldsymbol{k},-\boldsymbol{k}'}\delta_{j,j'}$	(S12)

Therefore, for any species, one obtains the following sum of acoustic and optical squared displacements ($j=a,o$):

$\left\langle\left( \xi_{\boldsymbol{n}i}- \xi_{\boldsymbol{0}\boldsymbol{i}} \right)^{2} \right\rangle\equiv{\delta R}_{\boldsymbol{n}i}^{2}=\frac{{4k}_{B}T}{Nm_{i}}\sum_{\boldsymbol{k}j} e_{i}^{j}\left( \boldsymbol{k} \right)e_{i}^{j}\left( \boldsymbol{-}\boldsymbol{k} \right)\frac{1-\cos(\boldsymbol{k}\cdot\boldsymbol{n})}{\Omega_{\boldsymbol{k}j}^{2}}$	(S13)

which can be summed over $i=1,2$ and integrated in the continuum $\boldsymbol{k}$-space. In the first case, the long-wavelength limit with $\Omega_{\boldsymbol{k}a}\approx lk\sqrt{\bar{\gamma}/m_{a}}$ still returns a random-walk law:

${\delta R}_{\boldsymbol{n}}^{2}\equiv\sum_{i} \frac{m_{i}}{m_{a}}{\delta R}_{\boldsymbol{n}i}^{2}\approx n\bar{a}^{2}$	(acoustic, $j=a$)	(S14)

the new step length $\bar{a}=\sqrt{n_{c}k_{B}T/\pi\bar{\gamma}}$ now depending on some average spring constant ($\bar{\gamma}$), with $n_{c}=2$. As the acoustic $m_{a}=\sum_{i} m_{i}$ denotes the total mass in the unit cell, Eq. (S14) is assigned the character of a probabilistic relation, where the individual displacements are weighted by means of single masses. Eq. (S14) is clearly affected by the number of atomic species in the unit cell. For two atomic species, it becomes:

${\delta R}_{\boldsymbol{n}}^{2} \equiv{\frac{m_{1}}{m_{1}+m_{2}}\delta R}_{\boldsymbol{n}1}^{2} + {\frac{m_{2}}{m_{1}+m_{2}}\delta R}_{\boldsymbol{n}2}^{2} \approx n\bar{a}^{2}$	(S15)

where $m_{1}$, $m_{2}$ are the masses of species $1$, $2$ and the step size $\bar{a}$ follows from mean $\bar{\gamma}$, averaged over the atomic interactions $11$, $22.$

For the faster vibrations of optical branches, it is convenient to integrate over the whole Brillouin’s zone and to approximate the dispersion relation to the mean value $\Omega_{\boldsymbol{k}o}\approx\sqrt{\bar{\gamma}} (1/\sqrt{m_{1}}+1/\sqrt{\mu})$, function of the reduced mass ($\mu$) and $m_{1}$, here supposed to be $\leq m_{2}$. In the end, Eq. (S14) returns a (roughly) constant contribution:

### ${\delta R}_{\boldsymbol{n}}^{2}\approx\frac{2\pi\bar{a}^{2}}{\bar{M}}$ (optical, $j=o$) (S15)

### where $\bar{M}=(m_{a}/m_{1}) {(1+ \sqrt{m_{1}/\mu})}^{2}$. The acoustic displacement will clearly prevail over the optical one in the limit of large $n$ values, i.e. $n\gg2\pi/\bar{M}$. An analytic inspection of this condition as a function of $x=m_{2}/m_{1}\geq1$ promptly reveals that $2\pi/\bar{M}$ is monotonically decreasing from the value $0.54$ ($x=1$) to zero ($x\to\infty$).

*Notes on Flory & Huggins’ (FH) solution theory*

The crucial quantity in the FH theory^18,19^ is the $\chi$ parameter:

$\chi=(z/2k_{B}T) (2u_{ps}-u_{pp}-u_{ss})$	(S16)

here defined for a homopolymer on a lattice with coordination number $z$. The mean free energies per site $u_{ps}$, $u_{pp}$, $u_{ss}$ quantify the interaction strength between individual units of polymer and solvent, polymer and polymer, solvent and solvent, respectively. We remind that pairwise interactions were demonstrated to be relevant to get the features of protein systems^20^.

The FH chi-parameter thus is basically a material–specific enthalpy term, depending on polymer–solvent interactions. Clearly, phase separation/aggregation is triggered by “poor” solvents^21^, or $\chi>0$. When the segregation degree $\chi N\gg1$ ($N=$polymer segment number), the enthalpy contribution to the separation largely prevails over the entropic effects, and a solution of (almost) pure phases in equilibrium forms^22^.

For a binary mixture of neutral molecules $A$,$B$ (e.g. a macromolecular blend), of arbitrary but fixed lengths with segment numbers $N_{A}$, $N_{B}$, the inflection point on the FH free energy curve returns a compositional-dependent critical point^23^ settled by:

$\phi_{Ac}= \sqrt{N_{B}}/(\sqrt{N_{A}}+\sqrt{N_{B}}$)	(S17)

${2\chi_{c}= (1/\sqrt{N_{A}}+1/\sqrt{N_{B}})}^{2}$	(S18)

where is the critical volume fraction ($\phi_{Ac}$) for the phase $A$.

The scattering function is known to depend as well on the single phase structure factors, the system composition ($\phi_{A}$,$\phi_{B}$) and interaction features ($\chi$)^24^. The more complex formal details for a polyphasic blend may be found elsewhere^25^. In this case, the conserved order parameter is a multicomponent vector gathering all the molar (or volume) fractions, whereas the FH parameter is mapped into a matrix like $\boldsymbol{\chi}_{ij}$.

*Notes on Associative Polymer Physics and Percolation*

Application of associative polymer physics to macromolecules of life has benefited from the growing consensus that a large number of membraneless biological condensates stem from the interplay of spontaneous or driven phase transformations with percolation phenomena^26^. Accordingly, a biological molecule is conceptualized as a series of stickers and spacers^27^. Stickers are recognized as elements responsible for attractive forces, while spacers, situated between stickers, are macromolecular segments that do not contribute significantly to attraction.

Consider, for simplicity, a phase transition between a dilute and a dense sticker phase, of concentrations $c_{-}$ and $c_{+}$. At some sticker concentration threshold ($c=c_{p}$), percolation will take place. It is a well-known networking transition dictated by the topology of connectivity points in the system. When $c_{-}\leq c_{p}<c_{+}$, phase separation implies percolation, and a mean field theory may be stated to evaluate $c_{p}$. For a mixture of different polymers with stickers $A$,$B$ it turns out that

${1/c}_{p}\approx\lambda_{AA}n_{A}^{2}+ \lambda_{AB}n_{A}n_{B}+\lambda_{BB}n_{B}^{2}$	(S19)

where $n_{k}$ denotes the number of stickers of the species $k$, and $\lambda_{ij}$ is a measure of the attractive volume related to the sticker pair $ij$^27^. The expression for the percolation threshold is easily extended to a sum over arbitrary sets of species and, as expected, is strongly composition-dependent.

**Table S1** C-Coord1 encoding.

| **C-Coord1** | A | C | D | E | F | G | H | I | K | L | M | N | P | Q | R | S | T | Y | V | W |
| --- | --- | --- | --- | --- | --- | --- | --- | --- | --- | --- | --- | --- | --- | --- | --- | --- | --- | --- | --- | --- |
| CA-Gly | 0 | 0 | 0 | 0 | 0 | 1 | 0 | 0 | 0 | 0 | 0 | 0 | 0 | 0 | 0 | 0 | 0 | 0 | 0 | 0 |
| Pro-MC | 0 | 0 | 0 | 0 | 0 | 0 | 0 | 0 | 0 | 0 | 0 | 0 | 2 | 0 | 0 | 0 | 0 | 0 | 0 | 0 |
| Carboxyl | 0 | 0 | 3 | 3 | 0 | 0 | 0 | 0 | 0 | 0 | 0 | 0 | 0 | 0 | 0 | 0 | 0 | 0 | 0 | 0 |
| Amide | 0 | 0 | 0 | 0 | 0 | 0 | 0 | 0 | 0 | 0 | 0 | 3 | 0 | 3 | 0 | 0 | 0 | 0 | 0 | 0 |
| His | 0 | 0 | 0 | 0 | 0 | 0 | 5 | 0 | 0 | 0 | 0 | 0 | 0 | 0 | 0 | 0 | 0 | 0 | 0 | 0 |
| Trp | 0 | 0 | 0 | 0 | 0 | 0 | 0 | 0 | 0 | 0 | 0 | 0 | 0 | 0 | 0 | 0 | 0 | 0 | 0 | 9 |
| Phe-Tyr | 0 | 0 | 0 | 0 | 6 | 0 | 0 | 0 | 0 | 0 | 0 | 0 | 0 | 0 | 0 | 0 | 0 | 6 | 0 | 0 |
| OH-Tyr | 0 | 0 | 0 | 0 | 0 | 0 | 0 | 0 | 0 | 0 | 0 | 0 | 0 | 0 | 0 | 0 | 0 | 1 | 0 | 0 |
| CH2 | 0 | 1 | 1 | 2 | 1 | 0 | 1 | 1 | 4 | 1 | 2 | 1 | 3 | 2 | 3 | 1 | 0 | 1 | 0 | 1 |
| CH | 0 | 0 | 0 | 0 | 0 | 0 | 0 | 1 | 0 | 1 | 0 | 0 | 0 | 0 | 0 | 0 | 1 | 0 | 1 | 0 |
| CH3 | 1 | 0 | 0 | 0 | 0 | 0 | 0 | 2 | 0 | 2 | 1 | 0 | 0 | 0 | 0 | 0 | 1 | 0 | 2 | 0 |
| OH | 0 | 0 | 0 | 0 | 0 | 0 | 0 | 0 | 0 | 0 | 0 | 0 | 0 | 0 | 0 | 1 | 1 | 0 | 0 | 0 |
| SH | 0 | 1 | 0 | 0 | 0 | 0 | 0 | 0 | 0 | 0 | 0 | 0 | 0 | 0 | 0 | 0 | 0 | 0 | 0 | 0 |
| S | 0 | 0 | 0 | 0 | 0 | 0 | 0 | 0 | 0 | 0 | 1 | 0 | 0 | 0 | 0 | 0 | 0 | 0 | 0 | 0 |
| NH3 | 0 | 0 | 0 | 0 | 0 | 0 | 0 | 0 | 1 | 0 | 0 | 0 | 0 | 0 | 0 | 0 | 0 | 0 | 0 | 0 |
| Arg | 0 | 0 | 0 | 0 | 0 | 0 | 0 | 0 | 0 | 0 | 0 | 0 | 0 | 0 | 4 | 0 | 0 | 0 | 0 | 0 |
| MC | 4 | 4 | 4 | 4 | 4 | 3 | 4 | 4 | 4 | 4 | 4 | 4 | 2 | 4 | 4 | 4 | 4 | 4 | 4 | 4 |

**Table S2** C-Coord2 encoding.

| **C-Coord2** | A | C | D | E | F | G | H | I | K | L | M | N | P | Q | R | S | T | Y | V | W |
| --- | --- | --- | --- | --- | --- | --- | --- | --- | --- | --- | --- | --- | --- | --- | --- | --- | --- | --- | --- | --- |
| Pro-MC | 0 | 0 | 0 | 0 | 0 | 0 | 0 | 0 | 0 | 0 | 0 | 0 | 1 | 0 | 0 | 0 | 0 | 0 | 0 | 0 |
| Carboxyl | 0 | 0 | 3 | 3 | 0 | 0 | 0 | 0 | 0 | 0 | 0 | 0 | 0 | 0 | 0 | 0 | 0 | 0 | 0 | 0 |
| Amide | 0 | 0 | 0 | 0 | 0 | 0 | 0 | 0 | 0 | 0 | 0 | 3 | 0 | 3 | 0 | 0 | 0 | 0 | 0 | 0 |
| His | 0 | 0 | 0 | 0 | 0 | 0 | 5 | 0 | 0 | 0 | 0 | 0 | 0 | 0 | 0 | 0 | 0 | 0 | 0 | 0 |
| Trp | 0 | 0 | 0 | 0 | 0 | 0 | 0 | 0 | 0 | 0 | 0 | 0 | 0 | 0 | 0 | 0 | 0 | 0 | 0 | 9 |
| Phe-Tyr | 0 | 0 | 0 | 0 | 6 | 0 | 0 | 0 | 0 | 0 | 0 | 0 | 0 | 0 | 0 | 0 | 0 | 6 | 0 | 0 |
| CH2 | 0 | 1 | 1 | 2 | 1 | 1 | 1 | 1 | 4 | 1 | 2 | 1 | 3 | 2 | 3 | 1 | 0 | 1 | 0 | 1 |
| CH | 0 | 0 | 0 | 0 | 0 | 0 | 0 | 1 | 0 | 1 | 0 | 0 | 0 | 0 | 0 | 0 | 1 | 0 | 1 | 0 |
| CH3 | 1 | 0 | 0 | 0 | 0 | 0 | 0 | 2 | 0 | 2 | 1 | 0 | 0 | 0 | 0 | 0 | 1 | 0 | 2 | 0 |
| OH | 0 | 0 | 0 | 0 | 0 | 0 | 0 | 0 | 0 | 0 | 0 | 0 | 0 | 0 | 0 | 1 | 1 | 1 | 0 | 0 |
| SH | 0 | 1 | 0 | 0 | 0 | 0 | 0 | 0 | 0 | 0 | 0 | 0 | 0 | 0 | 0 | 0 | 0 | 0 | 0 | 0 |
| S | 0 | 0 | 0 | 0 | 0 | 0 | 0 | 0 | 0 | 0 | 1 | 0 | 0 | 0 | 0 | 0 | 0 | 0 | 0 | 0 |
| NH3 | 0 | 0 | 0 | 0 | 0 | 0 | 0 | 0 | 1 | 0 | 0 | 0 | 0 | 0 | 0 | 0 | 0 | 0 | 0 | 0 |
| Arg | 0 | 0 | 0 | 0 | 0 | 0 | 0 | 0 | 0 | 0 | 0 | 0 | 0 | 0 | 4 | 0 | 0 | 0 | 0 | 0 |
| MC | 4 | 4 | 4 | 4 | 4 | 3 | 4 | 4 | 4 | 4 | 4 | 4 | 3 | 4 | 4 | 4 | 4 | 4 | 4 | 4 |

**Table S3** C-Coord3 encoding.

| **C-Coord3** | A | C | D | E | F | G | H | I | K | L | M | N | P | Q | R | S | T | Y | V | W |
| --- | --- | --- | --- | --- | --- | --- | --- | --- | --- | --- | --- | --- | --- | --- | --- | --- | --- | --- | --- | --- |
| Carbonyl-O | 0 | 0 | 2 | 2 | 0 | 0 | 0 | 0 | 0 | 0 | 0 | 1 | 0 | 1 | 0 | 0 | 0 | 0 | 0 | 0 |
| His | 0 | 0 | 0 | 0 | 0 | 0 | 5 | 0 | 0 | 0 | 0 | 0 | 0 | 0 | 0 | 0 | 0 | 0 | 0 | 0 |
| Trp | 0 | 0 | 0 | 0 | 0 | 0 | 0 | 0 | 0 | 0 | 0 | 0 | 0 | 0 | 0 | 0 | 0 | 0 | 0 | 9 |
| Phe-Tyr | 0 | 0 | 0 | 0 | 6 | 0 | 0 | 0 | 0 | 0 | 0 | 0 | 0 | 0 | 0 | 0 | 0 | 6 | 0 | 0 |
| CH2 | 0 | 1 | 1 | 2 | 1 | 1 | 1 | 1 | 4 | 1 | 2 | 1 | 3 | 2 | 3 | 1 | 0 | 1 | 0 | 1 |
| CH | 0 | 0 | 0 | 0 | 0 | 0 | 0 | 1 | 0 | 1 | 0 | 0 | 0 | 0 | 0 | 0 | 1 | 0 | 1 | 0 |
| CH3 | 1 | 0 | 0 | 0 | 0 | 0 | 0 | 2 | 0 | 2 | 1 | 0 | 0 | 0 | 0 | 0 | 1 | 0 | 2 | 0 |
| OH | 0 | 0 | 0 | 0 | 0 | 0 | 0 | 0 | 0 | 0 | 0 | 0 | 0 | 0 | 0 | 1 | 1 | 1 | 0 | 0 |
| SH | 0 | 1 | 0 | 0 | 0 | 0 | 0 | 0 | 0 | 0 | 0 | 0 | 0 | 0 | 0 | 0 | 0 | 0 | 0 | 0 |
| S | 0 | 0 | 0 | 0 | 0 | 0 | 0 | 0 | 0 | 0 | 1 | 0 | 0 | 0 | 0 | 0 | 0 | 0 | 0 | 0 |
| Nnoarom | 0 | 0 | 0 | 0 | 0 | 0 | 0 | 0 | 1 | 0 | 0 | 1 | 0 | 1 | 3 | 0 | 0 | 0 | 0 | 0 |
| C0 | 0 | 0 | 1 | 1 | 0 | 0 | 0 | 0 | 0 | 0 | 0 | 1 | 0 | 1 | 1 | 0 | 0 | 0 | 0 | 0 |
| MC | 4 | 4 | 4 | 4 | 4 | 3 | 4 | 4 | 4 | 4 | 4 | 4 | 4 | 4 | 4 | 4 | 4 | 4 | 4 | 4 |

**Table S4** C-Pos encoding.

| **C-Pos** | A | C | D | E | F | G | H | I | K | L | M | N | P | Q | R | S | T | Y | V | W |
| --- | --- | --- | --- | --- | --- | --- | --- | --- | --- | --- | --- | --- | --- | --- | --- | --- | --- | --- | --- | --- |
| CA-Gly | 0 | 0 | 0 | 0 | 0 | 1 | 0 | 0 | 0 | 0 | 0 | 0 | 0 | 0 | 0 | 0 | 0 | 0 | 0 | 0 |
| Pro-MC | 0 | 0 | 0 | 0 | 0 | 0 | 0 | 0 | 0 | 0 | 0 | 0 | 2 | 0 | 0 | 0 | 0 | 0 | 0 | 0 |
| CB | 1 | 1 | 1 | 1 | 1 | 0 | 1 | 1 | 1 | 1 | 1 | 1 | 0 | 1 | 1 | 1 | 1 | 1 | 1 | 1 |
| CB-Pro | 0 | 0 | 0 | 0 | 0 | 0 | 0 | 0 | 0 | 0 | 0 | 0 | 1 | 0 | 0 | 0 | 0 | 0 | 0 | 0 |
| Carboxyl | 0 | 0 | 3 | 3 | 0 | 0 | 0 | 0 | 0 | 0 | 0 | 0 | 0 | 0 | 0 | 0 | 0 | 0 | 0 | 0 |
| Amide | 0 | 0 | 0 | 0 | 0 | 0 | 0 | 0 | 0 | 0 | 0 | 3 | 0 | 3 | 0 | 0 | 0 | 0 | 0 | 0 |
| His | 0 | 0 | 0 | 0 | 0 | 0 | 5 | 0 | 0 | 0 | 0 | 0 | 0 | 0 | 0 | 0 | 0 | 0 | 0 | 0 |
| Trp | 0 | 0 | 0 | 0 | 0 | 0 | 0 | 0 | 0 | 0 | 0 | 0 | 0 | 0 | 0 | 0 | 0 | 0 | 0 | 9 |
| Phe-Tyr | 0 | 0 | 0 | 0 | 6 | 0 | 0 | 0 | 0 | 0 | 0 | 0 | 0 | 0 | 0 | 0 | 0 | 6 | 0 | 0 |
| OH-Tyr | 0 | 0 | 0 | 0 | 0 | 0 | 0 | 0 | 0 | 0 | 0 | 0 | 0 | 0 | 0 | 0 | 0 | 1 | 0 | 0 |
| CG | 0 | 0 | 0 | 1 | 0 | 0 | 0 | 2 | 1 | 1 | 1 | 0 | 1 | 1 | 1 | 0 | 1 | 0 | 2 | 0 |
| CD | 0 | 0 | 0 | 0 | 0 | 0 | 0 | 1 | 1 | 2 | 0 | 0 | 1 | 0 | 1 | 0 | 0 | 0 | 0 | 0 |
| CE | 0 | 0 | 0 | 0 | 0 | 0 | 0 | 0 | 1 | 0 | 1 | 0 | 0 | 0 | 0 | 0 | 0 | 0 | 0 | 0 |
| OH | 0 | 0 | 0 | 0 | 0 | 0 | 0 | 0 | 0 | 0 | 0 | 0 | 0 | 0 | 0 | 1 | 1 | 0 | 0 | 0 |
| SH | 0 | 1 | 0 | 0 | 0 | 0 | 0 | 0 | 0 | 0 | 0 | 0 | 0 | 0 | 0 | 0 | 0 | 0 | 0 | 0 |
| S | 0 | 0 | 0 | 0 | 0 | 0 | 0 | 0 | 0 | 0 | 1 | 0 | 0 | 0 | 0 | 0 | 0 | 0 | 0 | 0 |
| NH3 | 0 | 0 | 0 | 0 | 0 | 0 | 0 | 0 | 1 | 0 | 0 | 0 | 0 | 0 | 0 | 0 | 0 | 0 | 0 | 0 |
| Arg | 0 | 0 | 0 | 0 | 0 | 0 | 0 | 0 | 0 | 0 | 0 | 0 | 0 | 0 | 4 | 0 | 0 | 0 | 0 | 0 |
| MC | 4 | 4 | 4 | 4 | 4 | 3 | 4 | 4 | 4 | 4 | 4 | 4 | 2 | 4 | 4 | 4 | 4 | 4 | 4 | 4 |

**Table S5** AALetter encoding.

| **AALetter (One-hot)** | A | C | D | E | F | G | H | I | K | L | M | N | P | Q | R | S | T | Y | V | W |
| --- | --- | --- | --- | --- | --- | --- | --- | --- | --- | --- | --- | --- | --- | --- | --- | --- | --- | --- | --- | --- |
| A | 1 | 0 | 0 | 0 | 0 | 0 | 0 | 0 | 0 | 0 | 0 | 0 | 0 | 0 | 0 | 0 | 0 | 0 | 0 | 0 |
| C | 0 | 1 | 0 | 0 | 0 | 0 | 0 | 0 | 0 | 0 | 0 | 0 | 0 | 0 | 0 | 0 | 0 | 0 | 0 | 0 |
| D | 0 | 0 | 1 | 0 | 0 | 0 | 0 | 0 | 0 | 0 | 0 | 0 | 0 | 0 | 0 | 0 | 0 | 0 | 0 | 0 |
| E | 0 | 0 | 0 | 1 | 0 | 0 | 0 | 0 | 0 | 0 | 0 | 0 | 0 | 0 | 0 | 0 | 0 | 0 | 0 | 0 |
| F | 0 | 0 | 0 | 0 | 1 | 0 | 0 | 0 | 0 | 0 | 0 | 0 | 0 | 0 | 0 | 0 | 0 | 0 | 0 | 0 |
| G | 0 | 0 | 0 | 0 | 0 | 1 | 0 | 0 | 0 | 0 | 0 | 0 | 0 | 0 | 0 | 0 | 0 | 0 | 0 | 0 |
| H | 0 | 0 | 0 | 0 | 0 | 0 | 1 | 0 | 0 | 0 | 0 | 0 | 0 | 0 | 0 | 0 | 0 | 0 | 0 | 0 |
| I | 0 | 0 | 0 | 0 | 0 | 0 | 0 | 1 | 0 | 0 | 0 | 0 | 0 | 0 | 0 | 0 | 0 | 0 | 0 | 0 |
| K | 0 | 0 | 0 | 0 | 0 | 0 | 0 | 0 | 1 | 0 | 0 | 0 | 0 | 0 | 0 | 0 | 0 | 0 | 0 | 0 |
| L | 0 | 0 | 0 | 0 | 0 | 0 | 0 | 0 | 0 | 1 | 0 | 0 | 0 | 0 | 0 | 0 | 0 | 0 | 0 | 0 |
| M | 0 | 0 | 0 | 0 | 0 | 0 | 0 | 0 | 0 | 0 | 1 | 0 | 0 | 0 | 0 | 0 | 0 | 0 | 0 | 0 |
| N | 0 | 0 | 0 | 0 | 0 | 0 | 0 | 0 | 0 | 0 | 0 | 1 | 0 | 0 | 0 | 0 | 0 | 0 | 0 | 0 |
| P | 0 | 0 | 0 | 0 | 0 | 0 | 0 | 0 | 0 | 0 | 0 | 0 | 1 | 0 | 0 | 0 | 0 | 0 | 0 | 0 |
| Q | 0 | 0 | 0 | 0 | 0 | 0 | 0 | 0 | 0 | 0 | 0 | 0 | 0 | 1 | 0 | 0 | 0 | 0 | 0 | 0 |
| R | 0 | 0 | 0 | 0 | 0 | 0 | 0 | 0 | 0 | 0 | 0 | 0 | 0 | 0 | 1 | 0 | 0 | 0 | 0 | 0 |
| S | 0 | 0 | 0 | 0 | 0 | 0 | 0 | 0 | 0 | 0 | 0 | 0 | 0 | 0 | 0 | 1 | 0 | 0 | 0 | 0 |
| T | 0 | 0 | 0 | 0 | 0 | 0 | 0 | 0 | 0 | 0 | 0 | 0 | 0 | 0 | 0 | 0 | 1 | 0 | 0 | 0 |
| Y | 0 | 0 | 0 | 0 | 0 | 0 | 0 | 0 | 0 | 0 | 0 | 0 | 0 | 0 | 0 | 0 | 0 | 1 | 0 | 0 |
| V | 0 | 0 | 0 | 0 | 0 | 0 | 0 | 0 | 0 | 0 | 0 | 0 | 0 | 0 | 0 | 0 | 0 | 0 | 1 | 0 |
| W | 0 | 0 | 0 | 0 | 0 | 0 | 0 | 0 | 0 | 0 | 0 | 0 | 0 | 0 | 0 | 0 | 0 | 0 | 0 | 1 |

**Table S6** Chemical formula encoding.

| **Chemical formula** | A | C | D | E | F | G | H | I | K | L | M | N | P | Q | R | S | T | Y | V | W |
| --- | --- | --- | --- | --- | --- | --- | --- | --- | --- | --- | --- | --- | --- | --- | --- | --- | --- | --- | --- | --- |
| C | 3 | 3 | 4 | 5 | 9 | 2 | 6 | 6 | 6 | 6 | 5 | 4 | 5 | 5 | 6 | 3 | 4 | 9 | 5 | 11 |
| H | 5 | 5 | 4 | 6 | 9 | 3 | 8 | 11 | 13 | 11 | 9 | 6 | 7 | 8 | 13 | 5 | 7 | 9 | 9 | 10 |
| N | 1 | 1 | 1 | 1 | 1 | 1 | 3 | 1 | 2 | 1 | 1 | 2 | 1 | 2 | 4 | 1 | 1 | 1 | 1 | 2 |
| O | 1 | 1 | 3 | 3 | 1 | 1 | 1 | 1 | 1 | 1 | 1 | 2 | 1 | 2 | 1 | 2 | 2 | 2 | 1 | 1 |
| S | 0 | 1 | 0 | 0 | 0 | 0 | 0 | 0 | 0 | 0 | 1 | 0 | 0 | 0 | 0 | 0 | 0 | 0 | 0 | 0 |

**Table S7** Pos/Neg/HyPho encoding.

| **Pos/Neg/HyPho** | A | C | D | E | F | G | H | I | K | L | M | N | P | Q | R | S | T | Y | V | W |
| --- | --- | --- | --- | --- | --- | --- | --- | --- | --- | --- | --- | --- | --- | --- | --- | --- | --- | --- | --- | --- |
| Hydrophobicity | 1 | 0 | 0 | 0 | 1 | 0 | 0 | 1 | 0 | 1 | 1 | 0 | 0 | 0 | 0 | 0 | 0 | 1 | 1 | 1 |
| Positive | 0 | 0 | 0 | 0 | 0 | 0 | 1 | 0 | 1 | 0 | 0 | 0 | 0 | 0 | 1 | 0 | 0 | 0 | 0 | 0 |
| Negative | 0 | 0 | -1 | -1 | 0 | 0 | 0 | 0 | 0 | 0 | 0 | 0 | 0 | 0 | 0 | 0 | 0 | 0 | 0 | 0 |

**Table S8** NetChrg/SumHyPho encoding.

| **NetChrg/SumHyPho** | A | C | D | E | F | G | H | I | K | L | M | N | P | Q | R | S | T | Y | V | W |
| --- | --- | --- | --- | --- | --- | --- | --- | --- | --- | --- | --- | --- | --- | --- | --- | --- | --- | --- | --- | --- |
| Charge | 0 | 0 | -1 | -1 | 0 | 0 | 1 | 0 | 1 | 0 | 0 | 0 | 0 | 0 | 1 | 0 | 0 | 0 | 0 | 0 |
| Kyte-Doolittle | 1.8 | 2.5 | -3.5 | -3.5 | 2.8 | -0.4 | -3.2 | 4.5 | -3.9 | 3.8 | 1.9 | -3.5 | -1.6 | -3.5 | -4.5 | -0.8 | -0.7 | -1.3 | 4.2 | -0.9 |

**Table S9** Seq encoding.

| **Seq** | AAScan |
| --- | --- |
| A | 0.78 |
| C | 3.14 |
| D | 1.25 |
| E | 0.94 |
| F | 1.07 |
| G | 1.13 |
| H | 1.03 |
| I | 1.26 |
| K | 0.85 |
| L | 0.91 |
| M | 0.41 |
| N | 1.32 |
| P | 1.73 |
| Q | 0.93 |
| R | 1.75 |
| S | 1.31 |
| T | 1.57 |
| Y | 1.31 |
| V | 1.11 |
| W | 0.98 |

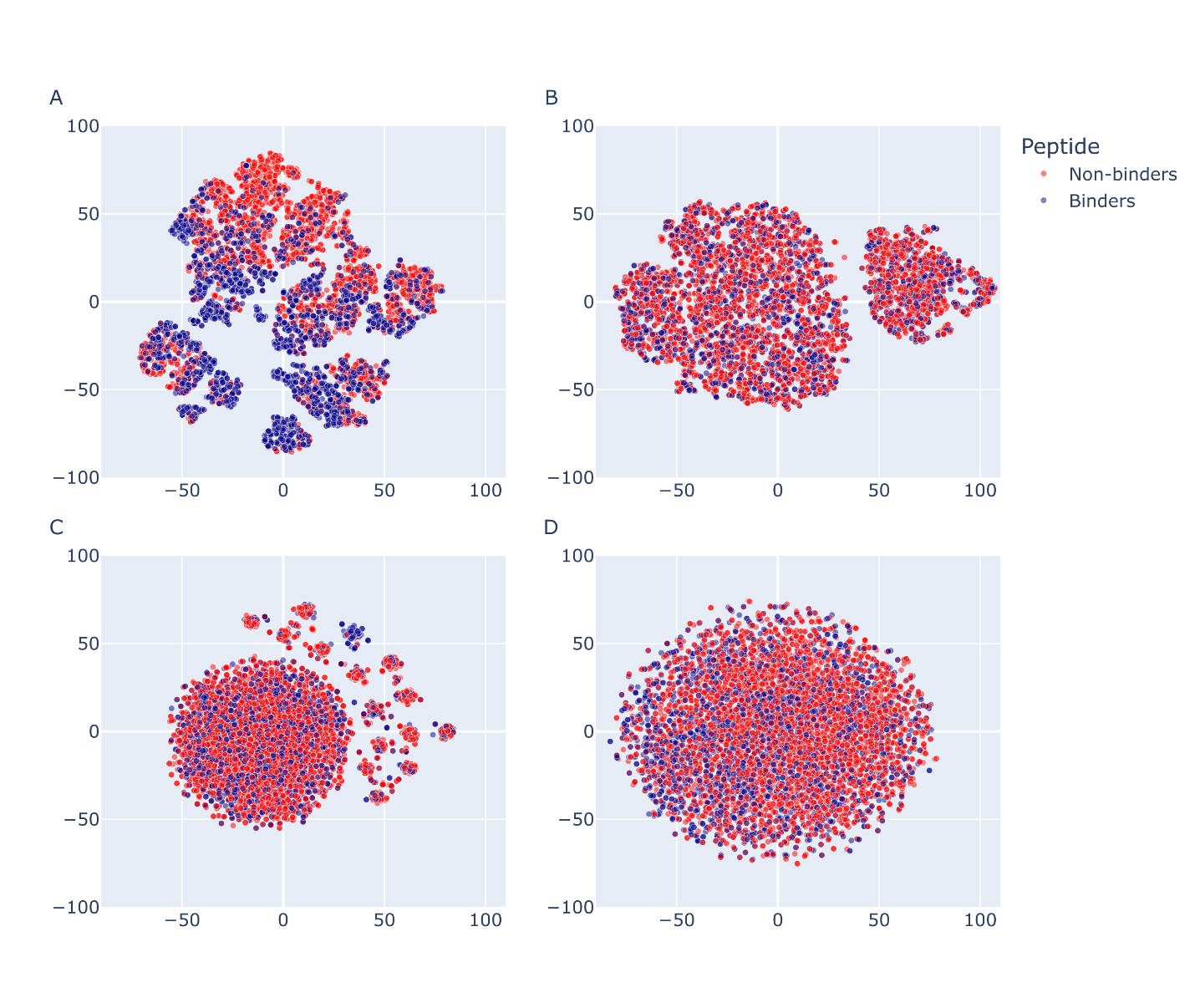
**Figure S1**. (A) t-SNE map showing microarray peptide distribution using atom type composition. The value assigned for each feature for each data point is the number of each atom type in a peptide sequence (C-Coord2), (B) FT-Seq representation of peptide sequence, (C) Seq representation of peptide sequence, and (D) UniSeq representation of peptide sequence. For all panels, red points signify binding peptides, and blue points signify non-binding peptides to survivin. The x and y axis signify the dissimilarity between peptides based on their respective representation. Perplexity value of 30.0 was used.

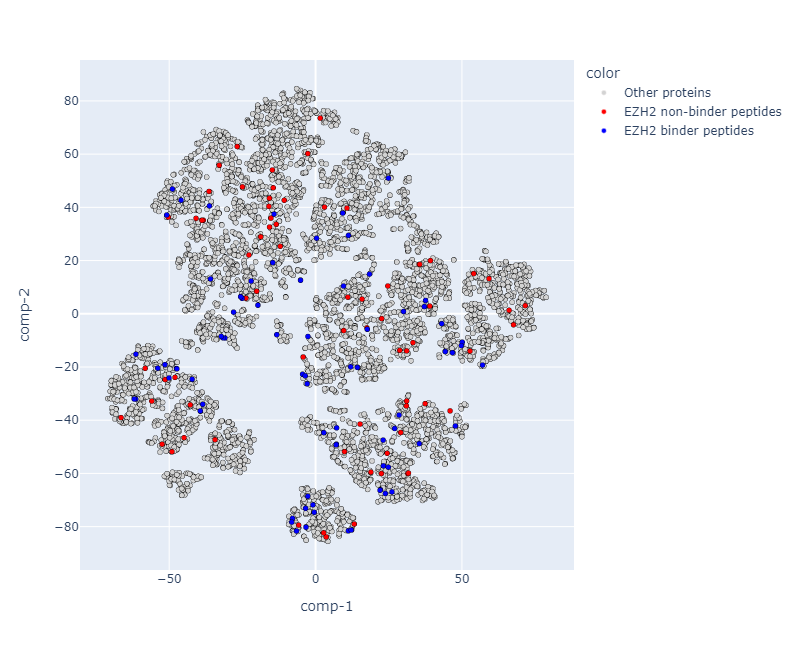

**Figure S2** t-SNE mapping of peptides from EZH2 subunit of PRC2. Red points signify binding peptides and blue points signify non-binding peptides to survivin. Peptides from other proteins are shown in grey. The x and y axis signify the dissimilarity between peptides based on their respective representation. Perplexity value of 30.0 was used.

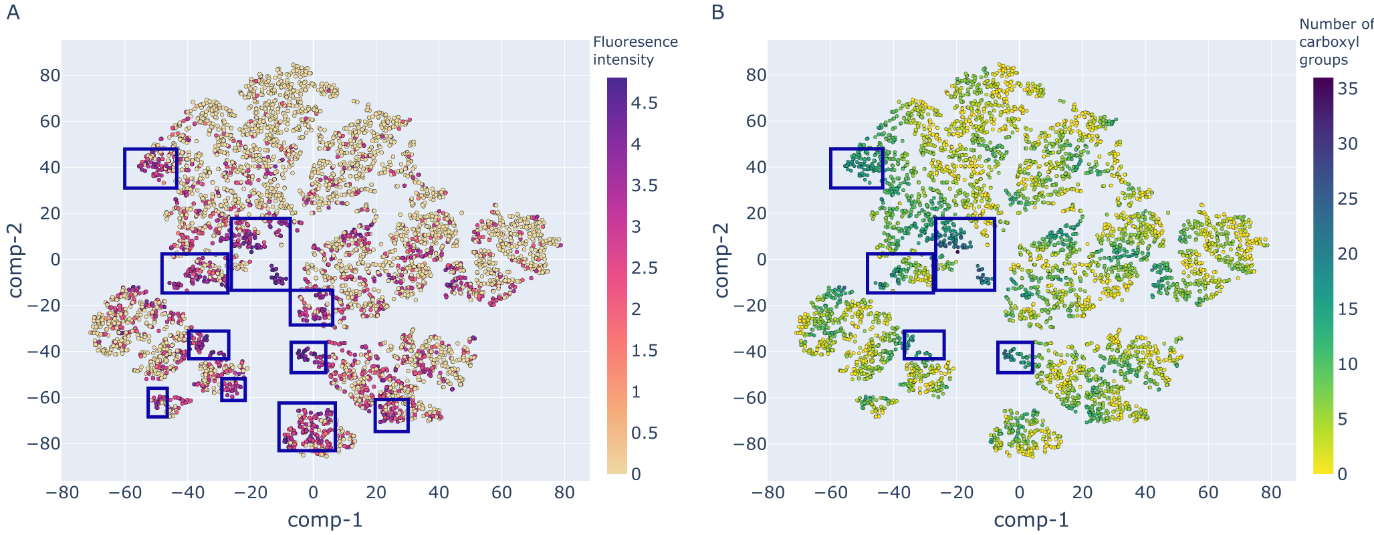
**Figure S3**. t-SNE distribution map using atom group composition of peptides (C-Coord2). (A) The colour ranges from 0 (non-binding) to 5 (the highest binding intensity), signifying fluorescence intensity spectrum across the peptide microarray dataset in a logarithmic scale. (B) The colour ranges from 0 (no atoms associated with carboxyl groups) to 35 (the highest number of atoms in carboxyl groups), signifying the number of carboxyl group in each peptide sequence. Perplexity value used was 30.0.

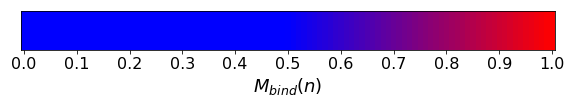

Aurora B

```
     70        80        90        100       110       120       130
     70 RHFTIDDFEIGRPLGKGKFGNVYLAREKKSHFIVALKVLFKSQIEKEGVEHQLRREIEIQAHLHHPNILR
     140       150       160       170       180       190       200
 140 LYNYFYDRRRIYLILEYAPRGELYKELQKSCTFDEQRTATIMEELADALMYCHGKKVIHRDIKPENLLLG
     210       220       230       240       250       260       270
 210 LKGELKIADFG---------------LDYLPPEMIEGRMHNEKVDLWCIGVLCYELLVGNPPFESASHNE
     281       291       301       311       321       331
 281 TYRRIVKVDLKFPASVPTGAQDLISKLLRHNPSERLPLAQVSAHPWVRANSRRVLPPS
```

Aurora A

```
     71        81        91        101       111       121       131
  71 -------------------------------------------------------RQWALEDFEIGRPLG
     141       151       161       171       181       191       201
 141 KGKFGNVYLAREKQSKFILALKVLFKAQLEKAGVEHQLRREVEIQSHLRHPNILRLYGYFHDATRVYLIL
     211       221       231       241       251       261       271
 211 EYAPLGTVYRELQKLSKFDEQRTATYITELANALSYCHSKRVIHRDIKPENLLLGSAGELKIADFGWSVH
     281       291       301       311       321       331       341
 281 APSSRRTTLAGTLDYLPPEMIEGRMHDEKVDLWSLGVLCYEFLVGKPPFEANTYQETYCRISRVEFTFPD
     351       361       371       381
 351 FVTEGARDLISRLLKHNPSQRPMLREVLEHPWITANSSK

```

STAT3

```
     71        81        91        101       111       121       131
  71 -----------------------------------------------------------------VVTEK
     141       151       161       171       181       191       201
 141 QQMLEQHLQDVRKRVQDLEQKMKVVENLQDDFDFNYKTLKSQGD---------SVTRQKMQQLEQMLTAL
     211       221       231       241       251       261       271
 211 DQMRRSIVSELAGLLSAMEYVQKTLTDEELADWKRRQQIACIGGPPNICLDRLENWITSLAESQLQTRQQ
     281       291       301       311       321       331       341
 281 IKKLEELQQKVSYKGDPIVQHRPMLEERIVELFRNLMKSAFVVERQPCMPMHPDRPLVIKTGVQFTTKVR
     351       361       371       381       391       401       411
 351 LLVKFPELNYQLKIKVCIDKDSGDVAALRGSRKFNILGTNTKVMNMEESNNGSLSAEFKHLTLREQRC—
     421       431       441       451       461       471       481
 421 -------ASLIVTEELHLITFETEVYHQGLKIDLETHSLPVVVISNICQMPNAWASILWYNMLTNNPKNV
     491       501       511       521       531       541       551
 491 NFFTKPPIGTWDQVAEVLSWQFSSTTKRGLSIEQLTTLAEKLLGP---YSGCQITWAKFCKENMAGKGFS
     561       571       581       591       601       611       621
 561 FWVWLDNIIDLVKKYILALWNEGYIMGFISKERERAILSTKPPGTFLLRFSESSKEGGVTFTWVE-----
     631       641       651       661       671       681       691
 631 --QIQSVEPYTKQQLNNMSFAEIIMGY--------LVSPLVYLYPDIPKEEAFGKYCR------------
     701       711
 701 –APYLKTKFICVTP
```

**Figure S4** Mapping of M_bind_(n) on the sequence of Aurora B, Aurora A and STAT3.
